# A novel bivalent interaction mode underlies a non-catalytic mechanism for Pin1-mediated Protein Kinase C regulation

**DOI:** 10.1101/2023.09.18.558341

**Authors:** Xiao-Ru Chen, Karuna Dixit, Yuan Yang, Mark I. McDermott, Hasan T. Imam, Vytas A. Bankaitis, Tatyana I. Igumenova

## Abstract

Regulated hydrolysis of the phosphoinositide phosphatidylinositol(4,5)-bis-phosphate to diacylglycerol and inositol-1,4,5-P_3_ defines a major eukaryotic pathway for translation of extracellular cues to intracellular signaling circuits. Members of the lipid-activated protein kinase C isoenzyme family (PKCs) play central roles in this signaling circuit. One of the regulatory mechanisms employed to downregulate stimulated PKC activity is via a proteasome-dependent degradation pathway that is potentiated by peptidyl-prolyl isomerase Pin1. Here, we show that contrary to prevailing models, Pin1 does not regulate conventional PKC isoforms α and βII via a canonical *cis-trans* isomerization of the peptidyl-prolyl bond. Rather, Pin1 acts as a PKC binding partner that controls PKC activity via sequestration of the C-terminal tail of the kinase. The high-resolution structure of Pin1 complexed to the C-terminal tail of PKCβII reveals that a novel bivalent interaction mode underlies the non-catalytic mode of Pin1 action.

Specifically, Pin1 adopts a compact conformation in which it engages two conserved phosphorylated PKC motifs, the turn motif and hydrophobic motif, the latter being a non-canonical Pin1-interacting element. The structural information, combined with the results of extensive binding studies and *in vivo* experiments suggest that non-catalytic mechanisms represent unappreciated modes of Pin1-mediated regulation of AGC kinases and other key enzymes/substrates.

**Impact statement:** Integrated biophysical, structural, and *in vivo* approaches demonstrate a non-canonical and non-isomerizable binding motif-dependent mode of protein kinase C regulation by the peptidyl-prolyl isomerase Pin1 in mammalian cells.

## INTRODUCTION

Protein Kinase C isoenzymes (PKCs) define a family of multi-modular Ser/Thr kinases that regulate cell growth, differentiation, apoptosis, and motility (1, 2). PKCs occupy a key node of the phosphoinositide signaling pathway whose intracellular arms are mediated by diacylglycerol (DAG) and IP_3_-dependent Ca^2+^ signaling (3). Novel and conventional PKCs are allosterically activated upon translocation to membranes. This membrane recruitment process involves PKC interactions with DAG and, in the case of Ca^2+^-dependent isoforms, the anionic phospholipids phosphatidylserine and PtdIns(4,5)P_2_ (4-6). Dysregulated PKC activity is implicated in cancer progression (7, 8), cardiac disease (9), diabetes (10), and neurodegenerative disorders (11). Both gain- and loss-of function PKC mutations are associated with diseased states, as reported recently for certain cancers (12, 13) and neurodegenerative disorders (14, 15).

Central to the question of PKC regulation is its phosphorylation state as it determines both cellular steady-state levels of the enzyme and its enzymatic activity. During the maturation process, a sequence of four ordered phosphorylation events (16) enable PKC to adopt a stable autoinhibited conformation in the cytosol. The autoinhibitory interactions between the N-terminal pseudo-substrate region and the C-terminal kinase domains are allosterically released by the interactions of second messengers with the PKC regulatory domain (**Fig. 1A**). The activated kinase can be additionally stabilized through the interactions with RACK (receptors for activated C-kinase) adaptor proteins (17, 18) that play an important role in the subcellular localization of PKCs (19). The open conformation that PKC assumes upon activation makes it susceptible to dephosphorylation by the PP2A (20) and PHLPP phosphatases (21, 22) with the result that the enzyme rapidly degraded in the cell. This “activation-induced” downregulation is one of the major mechanisms for terminating the PKC-mediated signaling response. Pin1, a peptidyl-prolyl isomerase, plays an important role in that process. It is for this reason that Pin1 was coined as the “molecular timer” for PKC lifetime (23).

**Figure 1.**
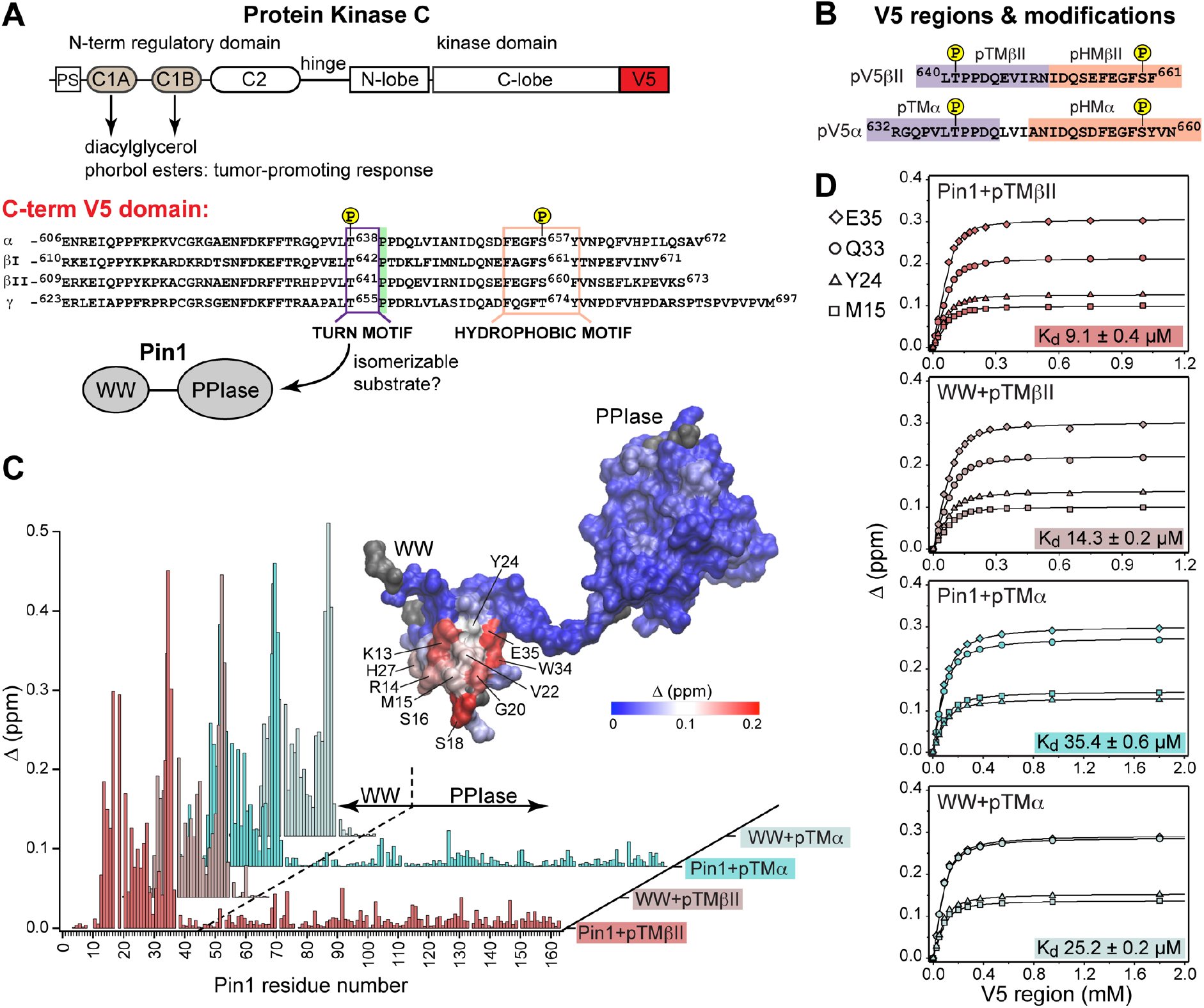
Pin1 binds the turn motifs of α and βII PKC isoenzymes through its WW domain. (**A**) Multi-modular architecture of conventional PKC isoenzymes, shown along with the amino acid sequence alignment of the C-terminal V5 domains. Turn and hydrophobic motifs are highlighted in purple and orange, respectively. (**B**) Notations for the V5 peptides that were selected for the NMR-detected binding experiments conducted in this study. (**C**) Residue-specific CSP plots of Pin1 and its isolated WW domain obtained upon binding of pTMβII and pTMα. The TM motifs interact exclusively with the WW domain. The CSP values Δ for the Pin1/ pTMβII complex are color-coded and mapped on to lowest-energy solution NMR structure of apo Pin1 (PDB: 1NMV). (**D**) Representative binding isotherms and dissociation constants for the formation of the complexes between turn motifs and Pin1 and its WW domain. The source 2D ^15^N-^1^H HSQC NMR spectra are given in **Figs. S1-S4**.

Pin1 is a peptidyl-prolyl isomerase of the parvuline family and is unique in its specificity towards pSer/pThr-Pro protein motifs (24). These Ser/Thr-Pro motifs constitute ∼30% of all phosphorylation sites in the proteome and their phosphorylation is catalyzed by proline-directed kinases (25). It is through isomerization of the *cis-trans* pSer/pThr-Pro bond that Pin1 brings the associated conformational changes of its substrates into biologically relevant timescales. In turn, conformation-specific activities of Pin1 client proteins are essential for the regulation of many cellular processes including metabolism, cell cycle progression, apoptosis, cell motility, cell proliferation, and cell survival (26). Pin1 is overexpressed/overactivated in cancers with the result that numerous oncoproteins are activated and tumor suppressor functions are deactivated (27-30). It is because Pin1 stimulates oncogenic pathways that Pin1 inhibitors show great promise in the development of new cancer therapies (31-37).

Pin1 consists of two domains – the WW and PPIase modules (38). The interplay between these domains is relevant to Pin1 catalytic function as it ensures the adaptability of Pin1 to a variety of phosphorylated substrates. Both Pin1 domains possess structural elements capable of interacting with pSer/pThr-Pro motifs, but only the isomerase domain has a catalytic role in the peptidyl-prolyl bond isomerization. The WW domain exhibits an affinity for Pin1 substrates that is ∼10-fold greater than that of the PPIase domain (39), and the WW domain is thought to either facilitate substrate recruitment to the PPIase active site and/or preferentially stabilize binding of a particular substrate isomer (40-42). The linker connecting the two domains confers significant flexibility to the Pin1 structure, and the conformational ensemble of Pin1 in solution is comprised of an ∼70:30 ratio of “compact” to “extended” conformers (43). Thus, Pin1 dynamics at both inter- and intra-domain levels play essential roles in the allosteric behavior of the enzyme (44, 45).

The current model of PKC regulation by Pin1 is based on extensive cell biological evidence. It posits that Pin1 catalyzes a *cis-trans* isomerization of the C-terminus of the conventional (or Ca^2+^-dependent) PKC isoforms α and βII (23). Herein, we report that, contrary to the prevailing model of Pin1 action, Pin1’s role in PKC α and βII regulation is a non-catalytic one. Instead, Pin1 acts as a PKC binding partner that sequesters two conserved phosphorylated PKC motifs in the disordered C-terminal tail (C-term) of the kinase. Our structure of Pin1-C-term of PKCβII complex reveals a novel bivalent interaction mode that has not been previously observed for any Pin1 complex. Our structural and biophysical data provide the molecular basis of non-catalytic Pin1 action that is also supported by the results of activity assays and *in vivo* experiments.

## RESULTS

### The turn motif of the α and βII PKC isoenzymes preferentially binds to the WW domain of Pin1

The key feature of the current model for Pin1-mediated PKC regulation is the Pin1-catalyzed *cis-trans* isomerization of the turn motif (TM) (**Fig. 1A**). TM is a conserved feature among many AGC kinases that presents a phosphorylatable Ser or Thr residue in the C-terminal domains of these enzymes (46). Phosphorylation of the TM is part of the PKC maturation process, and it is essential for enzyme stability (3, 47). In conventional PKC isoforms (α, βI/II, γ), the TM is followed by a proline residue and this pThr-Pro motif is proposed to be the isomerizable Pin1 target (41) (**Fig. 1A**). Therefore, the first step towards understanding how Pin1 regulates PKC was to determine the interaction mode between Pin1 and TM. To that end, NMR-detected binding experiments were conducted using uniformly ^15^N-enriched [U-^15^N] WW domain and full-length Pin1, and two synthetic peptides (pTMα and pTMβII) that correspond to the phosphorylated turn motifs of the α and βII PKC isoforms, respectively (**Fig. 1B**). Addition of increasing concentrations of pTMα and pTMβII resulted in drastic shifts in the 2D ^15^N-^1^H heteronuclear single-quantum coherence (HSQC) spectra of [U-^15^N] Pin1 and WW (**Figs. S1-S4**). The residue-specific chemical shift perturbation values (CSPs) were calculated for pairs of spectra corresponding to the maximum concentrations of pTM in each titration series *vs.* apo Pin1. The four CSP plots of **Fig. 1C** clearly demonstrate that the pTM binding site resides on the WW domain, and that the interaction mode is not significantly influenced by the presence of the PPIase domain in full-length Pin1. The fast exchange regime of the pTM-protein interactions enabled us to construct binding curves for all resolved residues with CSPs larger than the mean. The curves fit well with a single-site binding model (**Fig. 1D**). The reported K_d_ values fall into the micromolar range typical for WW-substrate affinities (42, 48, 49). The similarities of the K_d_ values between WW and full-length Pin1 (**Fig. 1D**) further support the conclusion that the PPIase domain does not significantly affect the pTM-WW interactions, and that the TMs of the α and βII PKC isoforms bind preferentially to the Pin1 WW domain.

### Pin1 does not catalyze the *cis*-*trans* isomerization of TM in α and βII PKC isoenzymes

The lack of high-affinity interactions of the PPIase domain with substrate does not preclude a catalytic mode of action for Pin1 as demonstrated previously for several Pin1 substrates (50). We therefore tested the ability of Pin1 to isomerize the TM. These experiments monitored activity against the isolated TM and in the context of a larger C-terminal region that harbors a second phosphorylated motif found in most AGC kinases – the hydrophobic motif (HM). Hydrophobic motifs in PKC isoforms are described by the sequence FXXF(<u>S/T</u>)(F/Y) where the underlined Ser/Thr is constitutively phosphorylated as part of the kinase maturation process (3, 51). HM is located downstream of the TM with 14 amino acid residues separating the two motifs. Cell biological studies suggest that the HM is involved in PKC interactions with Pin1 (23). However, the functional role of the HM is unclear as it does not fit the definition of a canonical Pin1 substrate due to the absence of a Pro residue that follows pSer/Thr. In these and all subsequent experiments, we used two synthetic double-phosphorylated peptides (pV5α and pV5βII; **Fig. 1B**) to model the C-term PKC regions that contain both phosphorylated HM and TM motifs.

Pin1 catalytic activity is readily measured using the ^1^H-^1^H exchange spectroscopy (EXSY) of substrates in the presence of catalytic amounts of the enzyme (52-54). Pin1 brings the slow *cis-trans* isomerization process of pSer/Thr-Pro bonds into biologically relevant timescales by enhancing the isomerization rate ∼10^3^-10^4^-fold. This rate enhancement gives rise to characteristic cross-peaks in the 2D ^1^H-^1^H EXSY NMR spectra. To our surprise, addition of catalytic amounts of Pin1 to either isolated pTMβII or to the entire phosphorylated pV5βII region failed to generate cross-peaks between the amide ^1^H resonances of pThr641 in the *trans-trans* (tt) and *cis-trans* (ct) conformations of the pThr641(−1)-Pro642(0)-Pro643(+1) segment (**Fig. 2A**). We intentionally used a long mixing time (0.5 s) to enable the detection of slow processes. Yet, no evidence of significant rate enhancement was observed. The lack of appreciable enhancement of the isomerization rate was similarly obtained for the pThr638(−1)-Pro639(0)-Pro640(+1) segment of the pTMα and pV5α regions (**Fig. 2B**). Of note, we were able to detect Pin1-catalyzed *cis-trans* isomerization of the peptidyl-prolyl bond between Gln634 and Pro635 in pTMα. This reaction was manifested by the appearance of cross-peaks between the amide ^1^H_N_ of Gln634 in the *cis* and *trans* conformations (**Fig. 2B**). Although Gln634-Pro635 is not a canonical Pin1 substrate, the catalytic action of Pin1 brings the reaction rate into a detectable range with k_EX_ of ∼1 s^-1^, where k_EX_ is the sum of the forward and reverse kinetic rate constants of the isomerization reaction, k_tc_ and k_ct_. The Gln634-Pro635 data serve as direct evidence that weak interactions of the turn motif region with the PPIase domain are sufficient for Pin1 to catalyze the *cis-trans* isomerization of the non-canonical Gln-Pro substrate but not the pThr-Pro segment of the turn motif. For both PKC α and βII isoforms, the HM has no detectable effect on Pin1 catalytic activity against these substrates.

**Figure 2.**
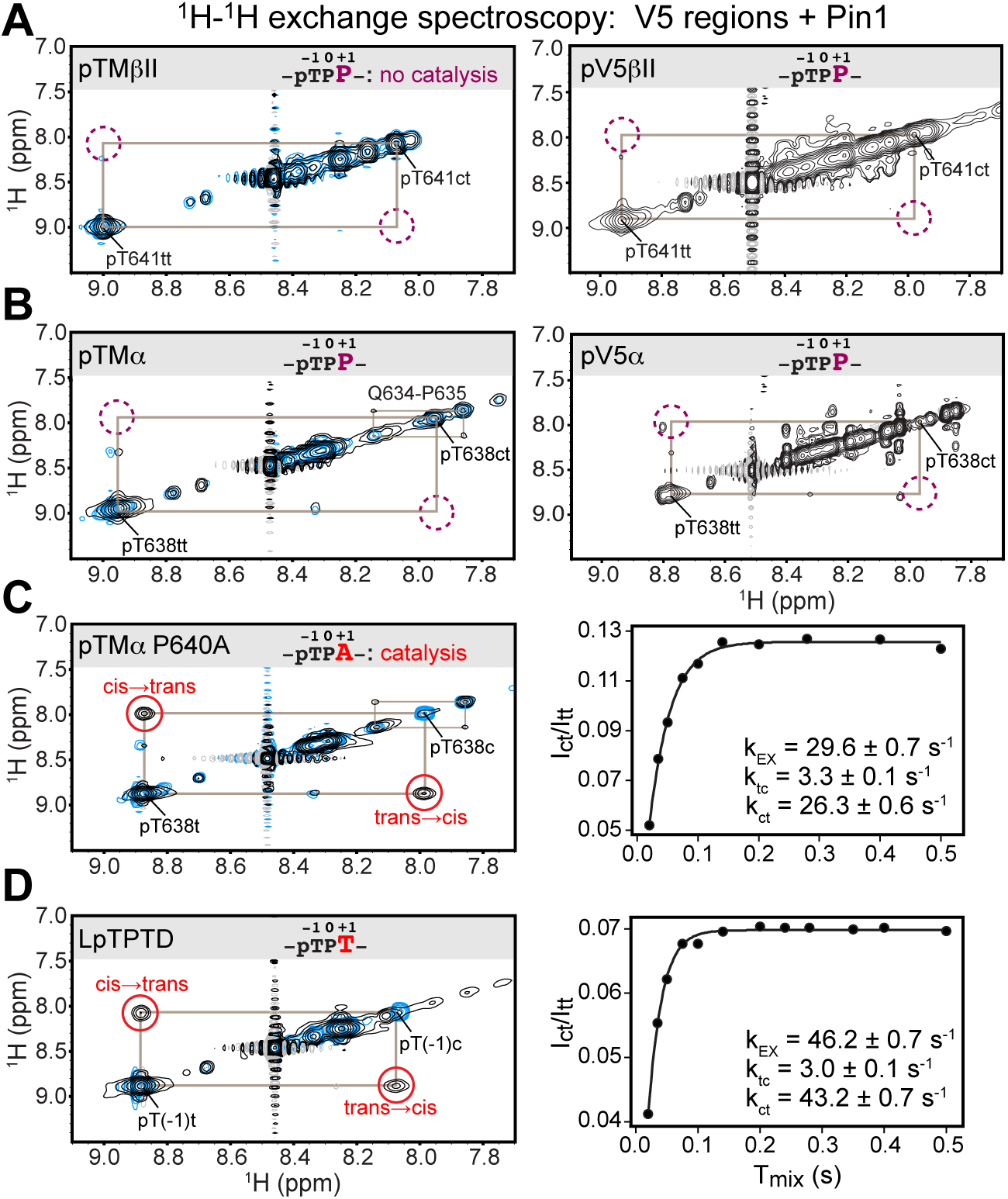
Pin1 does not appreciably catalyze isomerization of the turn motif in α and βII PKC isoenzymes due to presence of Proline at the +1 position. No exchange cross-peaks characteristic of Pin1-catalyzed pThr-Pro *cis-trans* isomerization are present in the spectra of TM regions from the PKC βII (**A**) and α (**B**) isoforms. This is demonstrated for both, isolated TM and the pV5 regions that harbor both hydrophobic and turn motifs. Non-specific catalysis by Pin1 is evident in the appearance of exchange cross-peaks for the ^1^H_N_ of Gln634 in *cis* and *trans* conformations of the Gln634-Pro635 segment (**B**). Replacement of Pro640 with Ala at the (+1) position of pTMα results in significant rate enhancement with the k_EX_ value of 29.6 s^-1^ (**C**). Thr at the (+1) position similarly enhances the rate of isomerization, as demonstrated for a short 5-residue LpTPPD peptide common to the TM regions of α, βII, and γ PKCs (**D**). The reference spectra collected with same parameters in the absence of Pin1 are shown in blue. The concentration of the V5 region peptides was 1-2 mM, with Pin1 added at catalytic amounts of 50 μM. The mixing times for all spectra are 0.5 s. The NMR spectra show the expansion of the ^1^H-^1^H amide region. Free peptide data (no Pin1 added) are shown in blue.

### The proline residue at the +1 position prohibits Pin1-mediated *cis*-*trans* isomerization of TM in α and βII PKC isoenzymes

We hypothesized that the proline at position +1 rendered the pTMα and pTMβII motifs poor substrates for PPIase-catalyzed isomerization. Thus, we generated the P640A variant of pTMα by replacing Pro at +1 position with Ala. Pin1-catalyzed isomerization of the pThr638-Pro639 bond in the P640A variant is readily detectable as reported by the appearance of distinct cross-peaks between the *cis* and *trans* conformers (**Fig. 2C**). The k_EX_ value of 29.6 s^-1^ obtained from the time dependence of the cross-peak intensities (see **Methods 4**) is typical for the Pin1 substrates. Similarly, replacement of Pro at the (+1) position with Thr in the short LpTPPD peptide common to the TM regions of α, βII, and γ PKC isoforms also resulted in significant enhancement of the *cis-trans* isomerization rate (k_EX_ value of 46.2 s^-1^; **Fig. 2D**). We conclude that Pro at the (+1) position of TM motifs is incompatible with these motifs serving as efficient substrates for isomerization by Pin1.

Our collective data further suggest that, contrary to prevailing models, Pin1 does not control downregulation of PKCα and βII isoforms by catalyzing isomerization of the PKC tail. Rather, Pin1 does so via a non-catalytic mechanism. The novelty of our findings prompted us to more thoroughly investigate the biophysical and structural basis of Pin1 interactions with the C-terminal domains of α and βII PKC isoforms. We focused on three aspects: (i) identifying the role of the hydrophobic motif, (ii) defining the binding mode, and (iii) establishing the effect of the C-terminal domain phosphorylation state on interactions with Pin1. To those ends, we conducted a total of 19 NMR-detected binding experiments between Pin1 and the relevant regions of the C-term domains of PKC α and βII isoforms with different phosphorylation states (**Table S1**). All information regarding the notations, sequences, and affinities is given in **Table S2**. The Pin1 domains are identified with the subscript “iso” and “Pin1” in their isolated forms and full-length Pin1 contexts, respectively.

### Hydrophobic motif of PKC binds to Pin1 via two independent sites

The hydrophobic motif FEGFpSF/Y (PKC α/βII) does not fit the canonical definition of a Pin1 substrate, and its role in the PKC-Pin1 interactions is unknown. Addition of the phosphorylated HM regions, pHMα and pHMβII to [U-^15^N] full-length Pin1 resulted in large CSPs of the amide N-H_N_ cross-peaks for many Pin1 residues, thereby providing direct and site-specific evidence for the Pin1-HM interactions (**Figs. S5A-B**). We were surprised to find that residues in both Pin1 domains were significantly affected, suggesting the existence of more than one pHM binding site in Pin1 (**Figs. 3A** and **S6A**). To identify which Pin1 domain harbors the HM binding site(s), we titrated isolated [U-^15^N] WW and PPIase domains with pHMβII and α (**Fig. 3B**, **S6B**). The CSP patterns of full-length Pin1 and the sum of isolated domains are essentially identical. These data indicate that Pin1 contains two sites that bind the phosphorylated hydrophobic motif: one residing in the WW domain and the other in the PPIase domain (**Figs. 3A,B** and **Fig. S6**).

**Figure 3.**
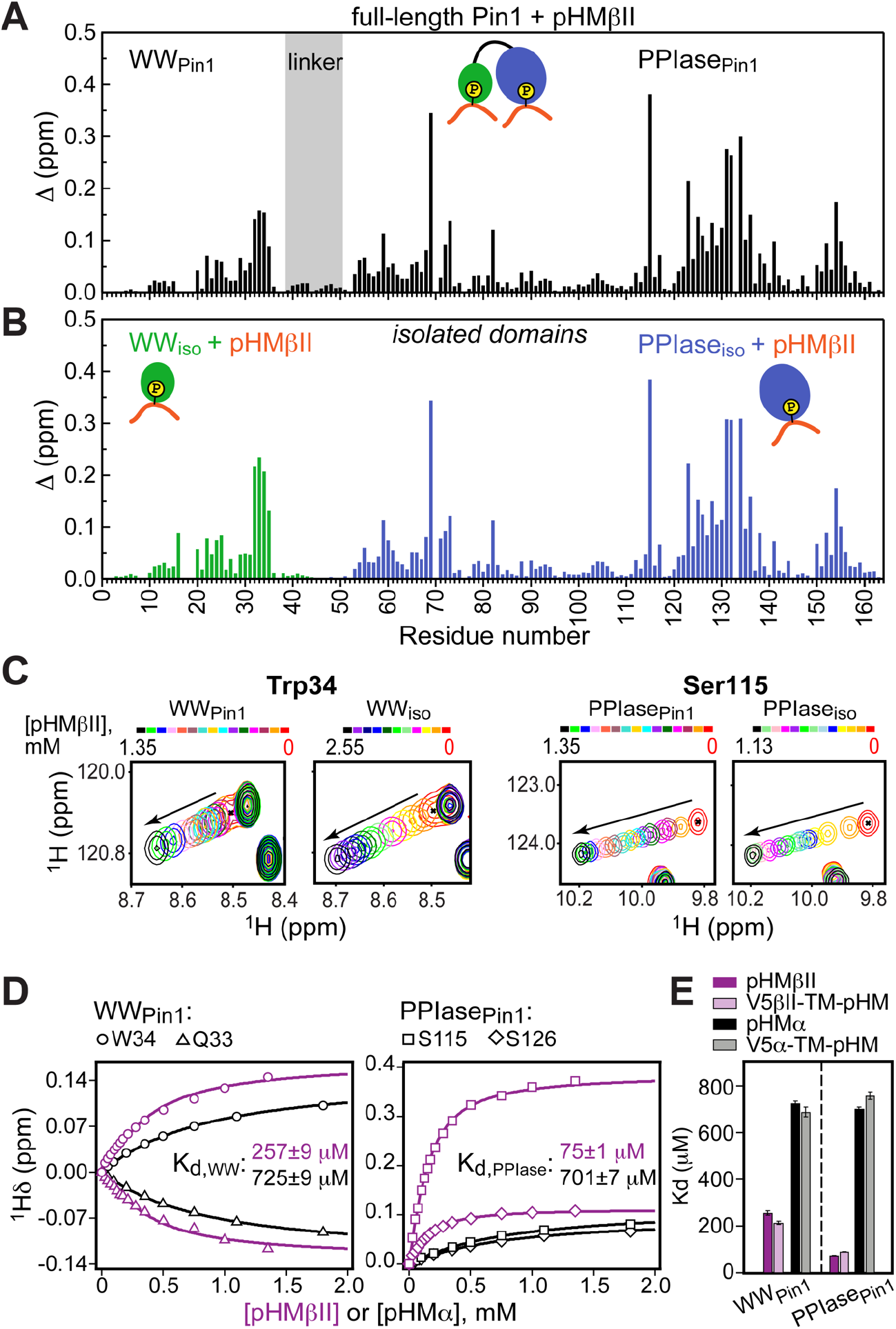
Hydrophobic motif interacts with Pin1 via two independent sites. Comparison of the CSP plots obtained at maximum concentrations of pHMβII versus ligand-free proteins for (**A**) full-length Pin1 and (**B**) isolated WW and PPIase domains. The similarity of the CSP patterns in (**A**) and (**B**) indicates that pHM has two independent binding sites in Pin1, one per domain. (**C**) Expansion of the ^15^N-^1^H chemical shift correlation spectra of Trp34 in the WW domain and Ser115 in the PPIase domain showing the fast-exchange regime of pHM binding. Full spectra are given in **Figure S5**. (**D**) Representative pHMβII (purple) and pHMα (black) binding curves for the Pin1 residues that belong to the WW and PPIase domains. Solid lines are the global fits to a model with 2 independent binding sites. (**E**) The unphosphorylated TM region upstream of the HM has no influence on the HM interaction mode with Pin1. Experimental conditions for all α and βII experiments are given in **Table S2**, experimental IDs # 5-10, 12, and 13.

The fast exchange regime of pHM binding to Pin1 (**Fig. 3C**) enabled us to construct the binding curves for all responsive residues and fit them globally using the two-site binding model (**Methods 3B**). The fitting produced domain-specific K_d_ values within full-length Pin1 (**Fig. 3D**). The pHMα motif has comparable affinities to both domains (725 μM to WW_Pin1_ and 701 μM to PPIase_Pin1_), whereas the pHMβII motif has a 3.5-fold greater affinity for PPIase_Pin1_ than it does for WW_Pin1_ (K_d_ values of 75 μM and 257 μM, respectively). To determine if the presence of the upstream unphosphorylated TM influences pHM-Pin1 interactions, we tested the binding of the V5α-TM-pHM and V5βII-TM-pHM regions to full-length Pin1. The NMR spectra and the measured K_d_ values are identical to those for the pHM regions only (**Fig. 3E**). Thus, unphosphorylated TM has little effect on the HM interactions with Pin1. The collective data demonstrate that, despite not being a canonical substrate, the phosphorylated hydrophobic motif of PKC isoenzymes can interact with Pin1 via two independent binding sites that reside on the WW and PPIase domains, respectively.

### Pin1 engages in unidirectional high-affinity bivalent interactions with the PKC C-terminus

Both the HM and TM motifs are phosphorylated in mature PKC. Addition of the pV5βII and pV5α regions that harbor both phosphorylated motifs to full-length Pin1 produced large CSP values in both Pin1 domains (**Figs. 4A**, **S7**, **S8A**). The CSP plot of full-length Pin1 complexed to pV5βII(α) matches the sum of the CSP plots obtained for the isolated WW domain complexed to pTMβII(α), and the isolated PPIase domain complexed to pHMβII(α) (**Figs. 4A,B; S7, S8**). These data provide unambiguous evidence that full-length Pin1 engages in bivalent interactions with the PKC C-terminal tail where WW binds the turn motif and PPIase binds the hydrophobic motif. The directionality of these interactions is imposed by the binding preferences of the turn motif. As the TM primarily interacts with the WW domain (**Fig. 1C**), the only site available for hydrophobic motif binding resides on the PPIase domain. Neither motif is an isomerizable Pin1 substrate.

**Figure 4.**
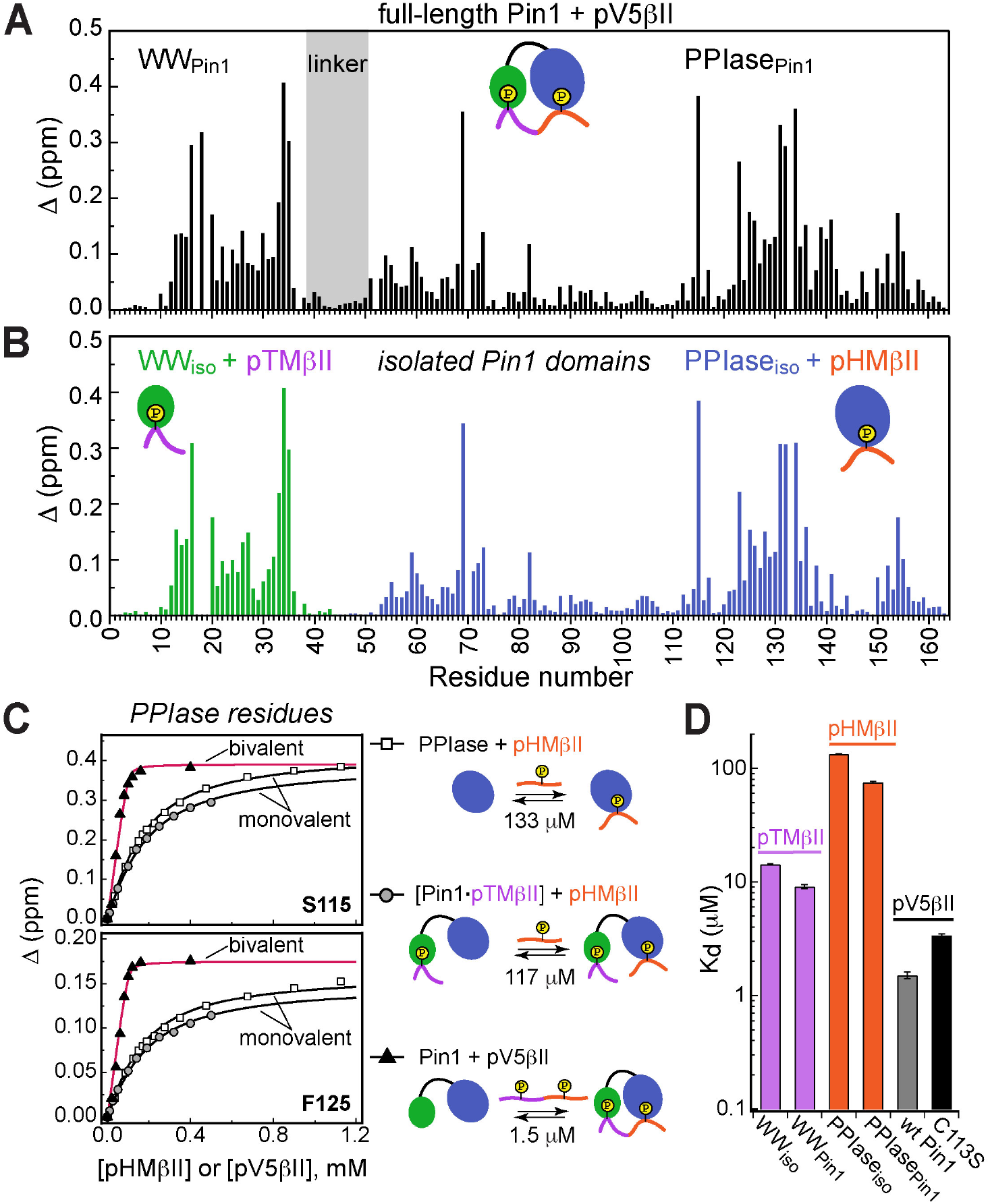
Unidirectional bivalent binding mode of the C-terminal PKCβII region to Pin1. (**A,B**) Comparison of the CSP plots of Pin1 obtained at maximum concentrations of pV5βII (**A**) and those of isolated domains, WW_iso_ and PPIase_iso,_ at maximum concentrations of pTMβII and pHMβII (**B**), respectively. The similarity of CSP patterns in (**A**) and (**B**) indicates that the C-term region of PKCβII binds to Pin1 in a unidirectional bivalent mode. (**C**) The TM and HM binding sites reside on the WW and PPIase domains, respectively. The protein concentration is 100 μM. Other details are given in **Table S2**, binding experiment IDs #9, 15, and 18. (**D**) K_d_ values for the monovalent interactions of the hydrophobic and turn motifs with isolated Pin1 domains and full-length Pin1 are contrasted with the K_d_ value for the bivalent Pin1-pV5βII interactions. ∼10-fold enhancement for the pTM binding to the WW domain and ∼90-fold enhancement of the pHM binding to the PPIase domain are attributed to bivalency. The K_d_ values used for this plot were obtained in the NMR-detected binding experiments. The K_d_ value for pV5βII binding to the catalytically deficient C113S Pin1 variant (black bar, 3.4 μM) exceeds the wild-type value by ∼2-fold.

Since the Pin1-pV5βII interactions fall into the intermediate-to-fast binding regime, NMR lineshape analysis was applied to obtain the K_d_ value of 1.5 μM (**Methods 3C**). These interactions are only moderately affected by the C113S mutation that reduces Pin1 catalytic activity ∼40-fold (55). The catalytically deficient C113S Pin1 variant shows a clear bivalent interaction mode with pV5βII (**Fig. S9, S10**). Compared to the wt Pin1, C113S has only ∼2-fold weaker affinity to pV5βII (K_d_ value of 3.4 μM, **Table S2**). The low micromolar K_d_ values obtained for WT Pin1 and the C113S variant reflect the thermodynamic advantage that the bivalent interaction mode imparts on the interactions with the C-term tail.

We illustrate the thermodynamic effect of bivalency using the binding curves of two representative residues (Ser115 and Phe125) in three distinct system compositions (**Fig. 4C**). The affinity of monovalent PPIase-pHMβII interactions does not appreciably depend on whether pTMβII is pre-bound to the WW domain: the corresponding K_d_ values are 133 μM and 117 μM, respectively (**Fig. 4C**). Moreover, the K_d_ values for the monovalent pHMβII-WW and pTMβII-PPIase interactions show little dependence on whether the isolated Pin1 domains or full-length Pin1 were used in the binding experiments (**Fig. 4D**). However, when both motifs are presented to full-length Pin1 on the same polypeptide chain, we observe an ∼90-fold enhancement in affinity (K_d_ = 1.5 μM). These general findings hold for the Pin1-pV5α interactions (**Fig. S11**). In aggregate, these data provide a complete thermodynamic view of bivalent interactions between Pin1 and the C-term PKC tail, and attribute the respective ∼10(3)-fold and ∼90(60)-fold enhancements of pTMβII(α) and pHMβII(α) binding to Pin1 to bivalency.

### Phosphorylation of the conserved motifs determines the Pin1-C-term interaction mode

The action of phosphatases on the HM and TM motifs generates monophosphorylated and dephosphorylated PKC species that are detected in Pin1 pull-down assays (23). To establish how these C-term modifications affect the Pin1 binding mode, we examined the interactions of Pin1 with the α and βII C-term regions where either TM (V5-pTM-HM), HM (V5-TM-pHM), or neither (V5) were phosphorylated. The CSP data, mapped onto the extended conformation of Pin1 for clarity (**Figs. 5A-D**), and the corresponding K_d_ values (**Figs. 3E, 5E**), revealed clear and consistent patterns.

**Figure 5.**
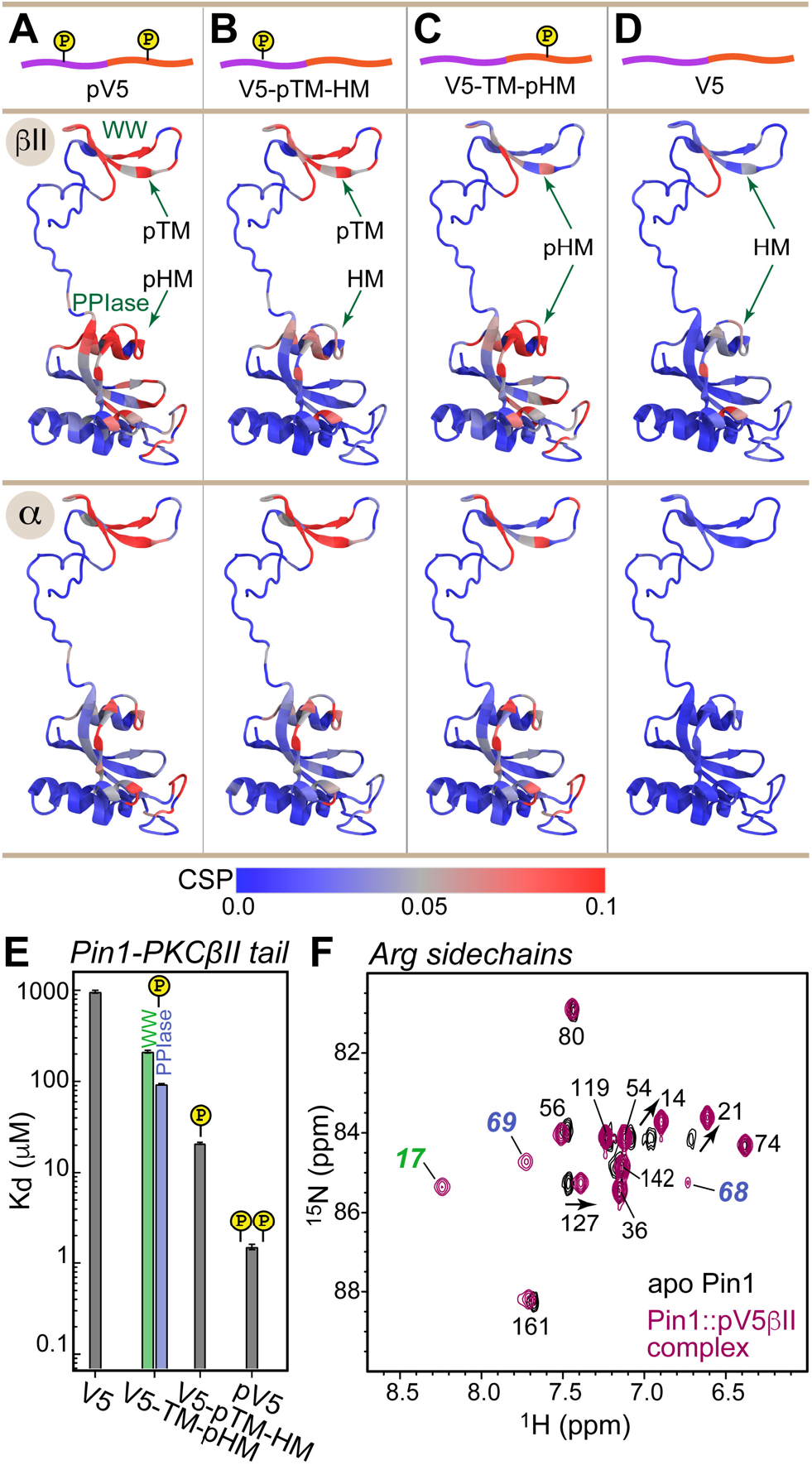
Phosphorylation state of the conserved C-term motifs defines the C-term interaction mode with Pin1. (**A-D**) CSP patterns of full-length Pin1 due to interactions with the C-term regions in different phosphorylation states. For clarity, the CSP values are mapped onto the extended NMR structure of apo Pin1 (1nmv). (**A,B**) Phosphorylation of pTM imposes a unidirectional bivalent mode irrespective of the phosphorylation state of the hydrophobic motif. The similarity of the Pin1 CSP patterns due to α and βII C-term binding suggest similar Pin1 interaction modes with PKCβII and PKCα isoforms. (**C**) Phosphorylation of HM directs hydrophobic motif to its binding sites on the WW and PPIase domains but does not impose a bivalent interaction mode. (**D**) Pin1 interactions with unphosphorylated C-term are only detectable for the C-term of PKCβII. (**E**) Dissociation constants of the Pin1::C-term(βII) complexes illustrating the enhancement of binding affinity due to phosphorylation. The data for the V5-TM-pHM binding to WW and PPIase are color-coded green and blue, respectively. The protein concentration is 100 μM. Other details are given in **Table S2**, binding experiment IDs #11, 13, 15, and 16. (**F**) 2D [^15^N-^1^H] HSQC spectra of the Arginine sidechains in apo Pin1 (black) and Pin1::pV5βII complex (maroon). Cross-peaks that are exchanged-broadened in apo Pin1 but reappear upon pV5βII binding belong to the Arg residues in the phosphate binding sites of WW (green) and PPIase (blue).

First, the C-term region singly phosphorylated at TM (V5-pTM-HM) produced a CSP pattern similar to that of the bivalent interaction between the fully phosphorylated pV5 region and Pin1 (**Figs. 5A,B; S12A-B**). Therefore, phosphorylated TM imposes a bivalent binding mode by employing its high-affinity interaction with the WW domain to direct the unphosphorylated HM to the binding site on the PPIase domain. Second, the C-term region singly phosphorylated at HM (V5-TM-pHM) produced a CSP pattern that is essentially identical to that of the isolated pHM binding to full-length Pin1 (**Figs. 5C**, **3A**, **S6A, S12C-D**). The unphosphorylated TM is unable to impose the bivalent binding mode with the result that the phosphorylated HM is free to occupy both the WW- and PPIase-localized binding sites. Third, the unphosphorylated C-term region interacts with Pin1 weakly. The CSPs are extremely small for the α isoform but are sufficiently large for βII to arrive at a K_d_ value of ∼960 μM (**Fig. 5D, S12E-F**). As the CSP pattern for the βII isoform is similar to that of the isolated pHM binding to the full-length Pin1 (**Fig. 3A, S12E-F**), we conclude that the phosphorylation status of TM determines the valency of the Pin1-PKC C-term interaction mode.

The K_d_ values determined for all C-term regions enabled us to estimate the thermodynamic gain associated with the phosphorylation of TM and HM in the context of bivalent interactions (**Fig. 5E** and **Table S2**). The affinity enhancement due to the TM phosphate is 62-fold, which corresponds to ΔΔG°_pTM_ of -2.4 kcal/mol. The affinity enhancement due to the HM phosphate is 14-fold, which corresponds to ΔΔG°_pHM_ of -1.6 kcal/mol. Both ΔΔG° values are within the range reported for the formation of the phosphate-mediated salt bridges in proteins (56). Summing up the contributions from the two phosphate groups produces the overall ΔΔG° of -4.0 kcal/mole, or ∼900-fold enhancement of the Pin1 affinity to the phosphorylated C-term of PKCβII.

The phosphate groups occupy the canonical phosphate binding sites of the Pin1 domains. We reach this conclusion by comparing the ^15^N-^1^H HSQC spectra of Arg sidechains in the apo and pV5βII-complexed Pin1 (**Fig. 5F**). Based on the structural data obtained previously for monovalent substrates (42, 57), Arg17_WW_ and the Arg68-Arg69_PPIase_ motif of the PPIase catalytic loop form salt bridges with the phosphate groups. While these three residues are exchanged-broadened in apo Pin1 due to dynamics, their cross-peaks reappear upon pV5βII binding -- consistent their direct interactions with these phosphate groups (**Fig. 5F**). To describe the bivalent interaction mode and identify the role of individual PKC and Pin1 residues, we determined the high-resolution structure of the complex.

### Structural basis of the Pin1-PKCβII C-term bivalent recognition mode

While both α and βII isoforms show similar bivalency patterns in their Pin1 interaction modes (**Fig. 5A-D**), the 8-fold higher binding affinity of pV5βII to Pin1 informed the choice of the Pin1-pV5βII complex for structural work. Since exhaustive screening of crystallization conditions failed to yield crystals suitable for X-ray diffraction, we pursued a solution NMR-based approach. The NMR structural ensemble of the Pin1-pV5βII complex was calculated with CYANA using ^1^H-^1^H NOEs, hydrogen bond and torsional angle restraints, and refined in explicit solvent using XPLOR-NIH (**Methods 5B**). The Pin1-pV5βII interface is defined by 75 inter-molecular NOEs whose assignment required the use of specifically labeled amino acids incorporated into the peptide substrate; a total of 9 different Pin1 complexes were prepared to ensure sufficient data redundancy (**Table S3**). The overall ensemble has a backbone and all-heavy-atom RMSD of 0.9 and 1.2 Å for the ordered regions, respectively (**Table S4** and **Fig. S13**).

The complex exhibits novel structural features that distinguish it from all other structures of Pin1 complexes known to date. First, the Pin1 substrate-binding mode is unusual in that pV5βII traverses the entire protein and interacts with both domains in a bivalent arrangement. The terminal pV5βII anchoring points are the two phosphorylated motifs, the pSer660 of the hydrophobic motif that binds to the PPIase domain and the pThr641 of the turn motif that binds to the WW domain (**Fig. 6A**). In addition, the pV5βII region in between the two phosphorylated motifs forms an extensive network of interactions with both WW and PPIase domains (**Figs. S14, S15**). The second noteworthy aspect of the structure is the extensive conformational change that Pin1 undergoes upon pV5βII binding. The WW and PPIase domains are brought into proximity primarily via interactions with the N-terminal pV5βII region. This compact conformation of pV5βII-complexed Pin1 is distinct from the “closed” conformation observed in available crystal structures of monovalent Pin1 complexes (**Fig. 6B**). The major differences are: (i) the ∼70° rotation of the WW domain relative to the PPIase module; and (ii) the repositioning of the α4 helix and the α4-β2 loop of the PPIase domain to accommodate pV5βII. The linker region connecting the WW and PPIase domains retains its flexibility and is not involved in pV5βII interactions.

**Figure 6.**
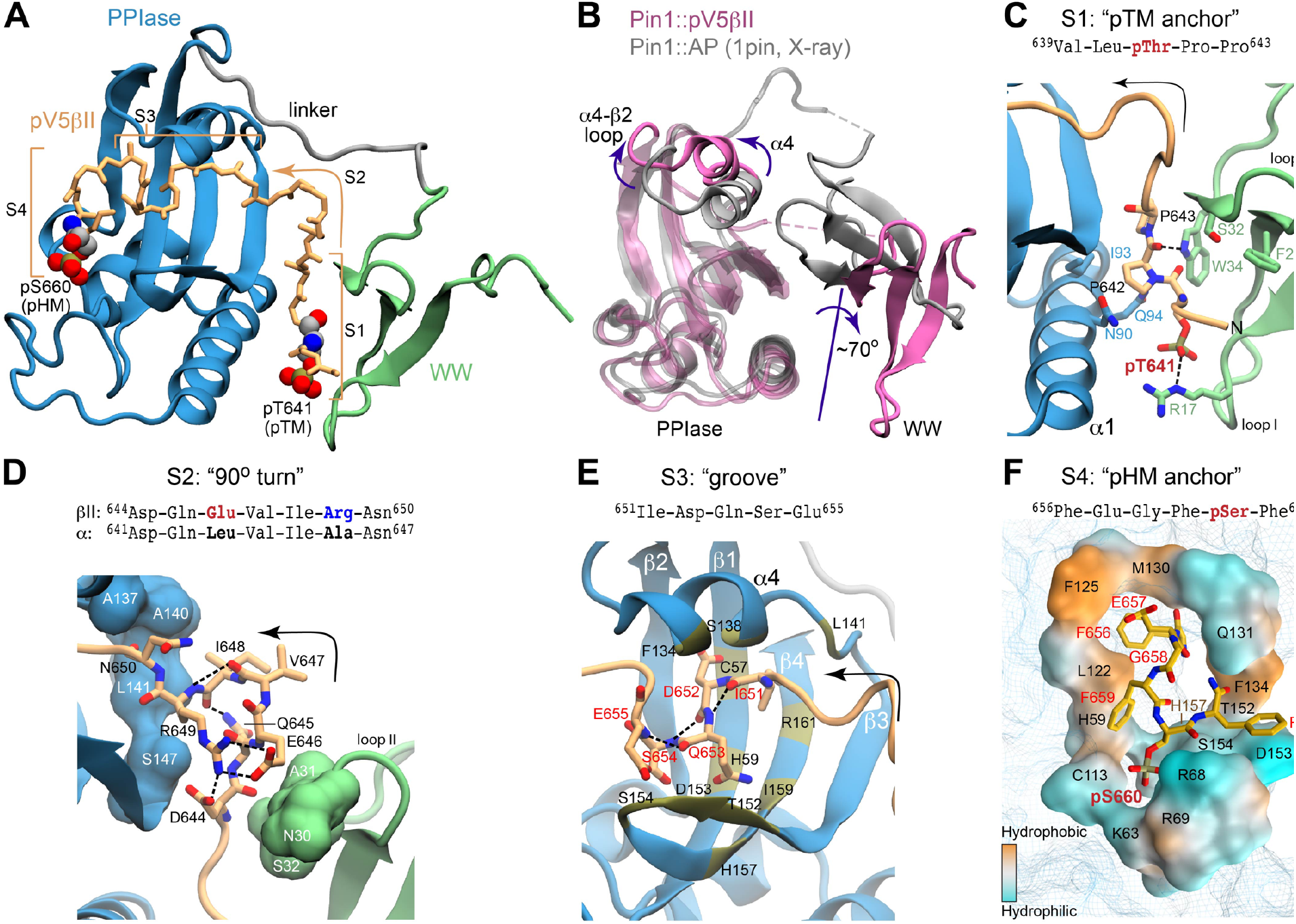
Structure of the Pin1::pV5βII complex reveals the bivalent recognition mode. (**A**) The lowest-energy NMR structure showing the pV5βII backbone (tan) forms an extensive binding interface with the WW (green) and PPIase (blue) domains of the full-length Pin1. pV5βII is broken into four segments, S1-S4, to facilitate the structural analysis. The phosphorylated Ser of the HM and Thr of the TM are shown in van der Waals representation. (B) Overlay of the crystal structure of the Pin1-AlaPro complex (1pin) and the NMR structure of the Pin1-pV5βII complex (8SG2, this work), illustrating the 70° rotation of the WW domain, along with the displacement of the α4 helix and the α4-β2 loop. Hydrogen bonds and salt bridges are shown with black dashed lines. (**C**) The “pTM anchor” segment is positioned at the interface between WW and PPIase domains. The phosphate group of pThr641 forms a salt bridge with Arg17. (**D**) The “90 turn” segment is stabilized by intramolecular hydrogen bonds and is wedged between the WW and PPIase domains. (**E**) The “groove” segment is threaded between the α4 helix and the β3-β4 hairpin of PPIase. (**F**) The “pHM anchor” segment occupies the catalytic site of the PPIase domain. Residues forming the site are color-coded according to amphiphilicity. The phosphate group of pSer660 forms salt bridges with the Arg68 and Arg69 residues of the catalytic loop.

### Interface of the pTM anchor and turn regions of PKCβII C-term with Pin1

To facilitate structural analysis of the Pin1-pV5βII interface, we separated pV5βII into four segments: the “pTM anchor” (639-643, S1), the “90° turn” (644-650, S2), the “groove” (651-655, S3), and the “pHM anchor” (656-661, S4) (**Fig. 6A**). The 5-residue “pTM anchor” harboring the pThr641-Pro642-Pro643 segment is positioned at the interface between the WW and PPIase domains (**Fig. 6C**). Its backbone runs almost parallel to the α1 helix of the PPIase domain. The key interactions are the salt bridge between the Thr641 phosphate and guanidinium groups of Arg17_WW_, and the hydrophobic contacts of the pThr641 methyl group and Pro642 pyrrolidine ring with the sidechains of Trp34_WW_, Asn90_PPIase_, Ile93_PPIase_ and Gln94_PPIase_. In addition, the NHε group of the Trp34_WW_ forms a hydrogen bond with the carbonyl oxygen of pThr641.

The Pro643-Asp644 segment reorients the backbone such that it runs along the β1 strand of the WW domain. This marks the beginning of the 7-residue “90° turn” segment (**Fig. 6D**) that completes the inter-domain arm of pV5βII and is responsible for the major realignment of the PPIase-WW domain interface. Prior structural work on Pin1 identified residues 137-141_PPIase_, 148-149_PPIase_, and 28-32_WW_ as being involved in the dynamic inter-domain interface (48, 49). In the Pin1-pV5βII complex, many residues from this subset are now engaged in interactions with “90° turn” segment pV5βII (**Fig. 6D** and **Fig. S14**). The turn itself is stabilized by intra-pV5βII hydrogen bonds and salt bridges. Arg649 plays a particularly prominent role in that regard as it participates in both types of interactions. The intra-molecular salt bridge formed by the Arg649 and Glu646 sidechains is unique to the βII isoform (**Fig. S16**) because these Arg and Glu residues are replaced by hydrophobic Leu and Ala/Met residues in other conventional PKC isoforms (**Fig. 6D** and **Fig. 1A**). The contributions of Arg sidechain-mediated interactions are likely responsible for the ∼8-fold higher affinity of pV5βII for Pin1 relative to the affinity of pV5α.

### Interface of the groove and pHM anchor regions of PKCβII C-term with Pin1

The turn is followed by the 5-residue pV5βII “groove” segment that is rich in polar residues and interacts exclusively with the PPIase domain. The groove segment threads through a deep groove in the Pin1 PPIase domain formed by helix α4 and the β-sheet comprised of strands β3, β4, and β1 (**Fig. 6E** and **S17**). This configuration drives the repositioning of helix α4 and the α4-β2 loop relative to the structures of Pin1 complexed to the monovalent substrate (**Fig. 6B**). The Pin1-pV5βII interface involves hydrophobic contacts of the only hydrophobic residue of this pV5βII segment (Ile651) with Pin1 residues Leu141_PPIase_, I159_PPIase_ and the Arg161_PPIase_ methylenes. Among polar residues, pV5βII residues Ser654 and Glu655 are within H-bonding distance with the three Pin1 residues of the conserved parvulin tetrad (58-60): His59_PPIase_ and His157_PPIase_, and Thr152_PPIase_, respectively (**Fig. S15**).

The last segment of pV5βII (the “pHM anchor”) emerges from the PPIase groove and occupies the catalytic site. This “anchor” segment contains the entire PKCβII hydrophobic motif whose key residues (Phe656, Phe659, and pSer660) interact with the amphiphilic environment of the Pin1 catalytic site (**Fig. 6F**). Specifically, the sidechains of Phe656 and Phe659 are accommodated by the hydrophobic environment formed by Met130_PPIase_, Phe125_PPIase_, and Leu122_PPIase_. The Phe659 aromatic ring also engages in stacking interactions with His59_PPIase_.

The pSer660 phosphate is anchored to the Pin1 catalytic loop via salt bridges with a triad of positively charged residues (Arg68_PPIase_, Arg69 _PPIase_, and Lys63 _PPIase_). Those interactions rigidify the loop as is evident from the reappearance of Arg68 _PPIase_ and Arg69 _PPIase_ resonances in the NMR spectra upon complex formation (**Fig. 5F**). Phe661, the third Phe of the HM, is not involved in any persistent interactions with the Pin1 PPIase module.

We then compared the binding poses of pHM with that of the D-peptide -- a potent unnatural peptide inhibitor of Pin1 that binds specifically to the catalytic site of the PPIase domain (61) (**Figs. S18A-B**). Structural overlay of the complexes shows that pHM and the D-peptide occupy the same PPIase region, and the positions of the ligand phosphate groups coincide (**Fig. S18C**). Notable differences include the arrangement of the ligand hydrophobic ring moieties in the catalytic site, and the position of the Gln131_PPIase_ sidechain relative to the ligand. Specifically, the space occupied by the Gln131_PPIase_ sidechain in the Pin1::pV5βII complex is occupied by the Gln5 sidechain of the D-peptide in the Pin1::D-peptide complex (**Fig. S18C**). Gln131 is the C-terminal residue of the α4 helix that undergoes the most significant rearrangement upon the formation of the bivalent Pin1::pV5βII complex (**Fig. 6B**).

### Pin1 null HEK293T cells exhibit elevated steady-state levels of PKCα

The structural data revealed: (i) a bivalent mode of Pin1 interactions with the C-term tail of PKCβII, (ii) that the phosphate groups occupy the canonical binding sites of the Pin1 WW and PPIase domains, and (iii) that the intervening residues form an extensive network of interactions with the residues of both Pin1 domains. Moreover, the similarities of the NMR chemical shift perturbation patterns between pV5βII and pV5α binding to Pin1 report that the bivalent interaction mode is shared by both the α and βII PKC isoforms. To interrogate our biophysical and structural conclusions regarding Pin1 function in an *in vivo* context, we developed a system for assessing Pin1-mediated PKCα regulation without the contribution of endogenous Pin1 activity in a HEK293T cell model. PKCα was chosen for these analyses because we were able to detect endogenous PKCα by immunoblotting in HEK293T cells, whereas PKCβII over-expression was required for detection in this cell line.

Previous studies reported Pin1 downregulates PKC levels in serum-starved cells stimulated with PDBu (23). Consistent with this general concept, efficient (∼80%) siRNA-mediated knockdown of Pin1 expression resulted in an ∼45% elevation in steady-state PKCα levels in cells cultured in the presence of serum. That is, cells stimulated by natural agonists under more physiological conditions than those stimulated by exogenous agonist PDBu under serum starvation (**Fig. S19A**). This relationship was further corroborated by CRISPR screens that produced Pin1 null HEK293T cell lines (**Methods 6F; Fig. S19B**). Clonally-derived cells recovered from the CRISPR challenge exhibited a wide range of Pin1 expression levels, and immunoblotting confirmed an inverse correlation between endogenous Pin1 and steady-state PKCα levels (**Fig. 7A**; R = -0.92, p<0.0005).

**Figure 7.**
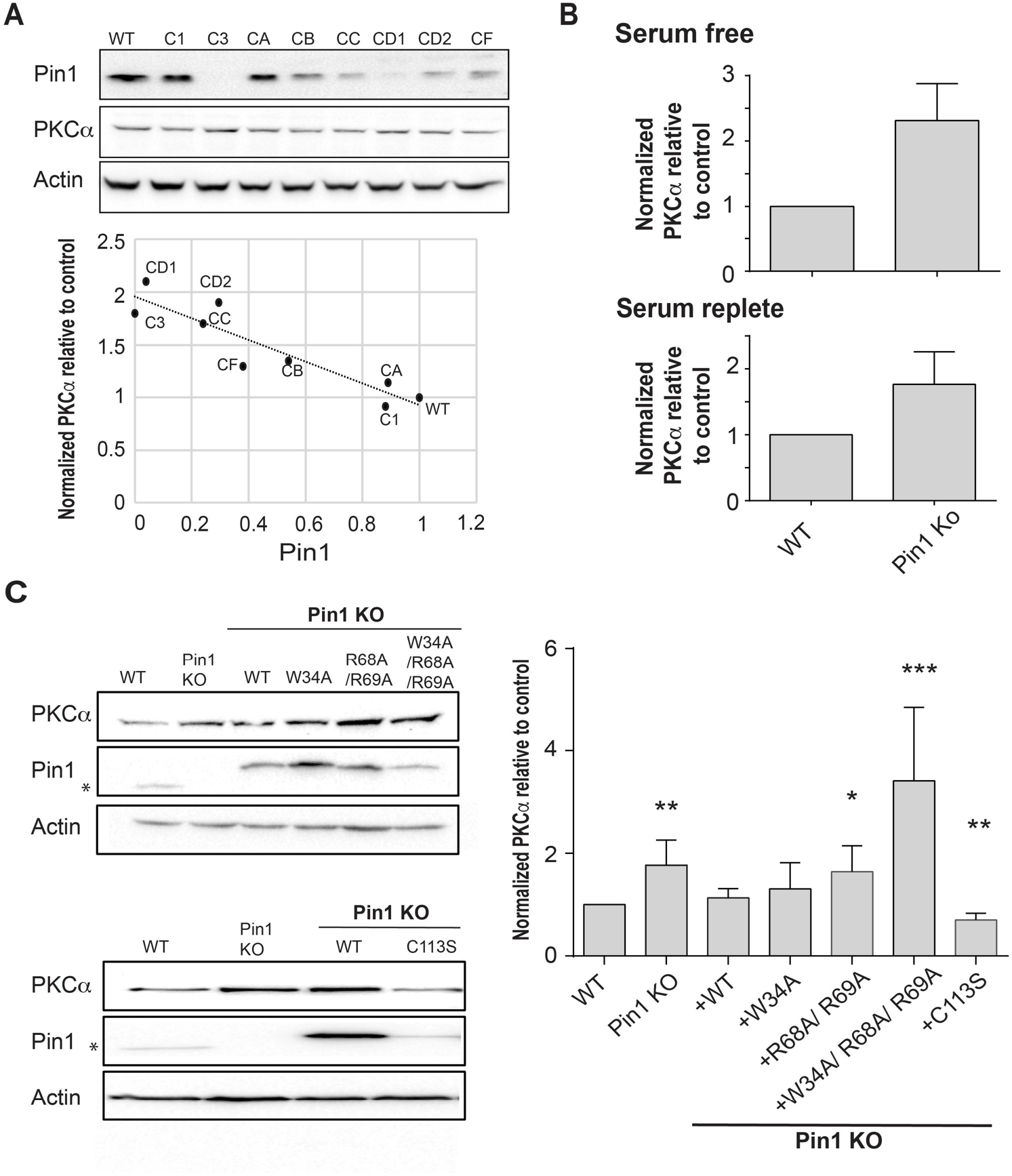
Regulation of PKCα homeostasis by Pin1 in HEK293T cells. (**A**) PKCα protein levels at steady-state are inversely proportional to Pin1 levels. HEK293T cells were transfected with CRISPR/Cas9 plasmids encoding Pin1 guide RNAs and clonal lines were generated. Lysates of the clonally-derived cell lines were resolved by SDS-PAGE, transferred to nitrocellulose, and immunoblots developed to visualize PKCα, Pin1 and actin. Blot profiles are shown at top. Bottom panel relates steady-state PKCα protein levels to Pin1 steady-state levels. Actin was used to normalize the PKCα and Pin1 profiles for each cell line and the PKCα/actin ratio was set as 1.0 for parental WT Pin1 cells. Clone 3 (C3) expressed no detectable Pin1 antigen and was selected for further study. (**B**) Pin1 null HEK293T cells exhibit elevated steady-state PKCα protein levels when cells are incubated under serum-free (upper panel; 2.3 ± 0.2; n=7; p < 0.0001, two-tailed t-test) and serum-replete conditions. (lower panel; 2.3 ± 0.3; n=12; p < 0.0002, two-tailed t-test). Actin was used to normalize the PKCα and Pin1 immunoblot profiles for each cell line and the PKCα/actin ratios were set as 1.0 for parental WT Pin1 cells. **(C)** PKCα regulation by Pin1 derivatives with defined biochemical defects. Left panels: At top are shown representative PKCα, Pin1 and actin immunoblot profiles for WT HEK293T cells, Pin1 KO cells (C3) and Pin1 KO cells stably expressing the indicated mutant Pin1 proteins defective in PKCα binding. At bottom are shown representative PKCα. Pin1 and actin immunoblot profiles for WT HEK293T cells, Pin1 KO cells (C3) and Pin1 KO cells stably expressing the ‘catalytic-dead’ Pin1^C113S^ mutant. In both panels the asterisk denotes endogenous Pin1 as the ectopically expressed Pin1 proteins are 3.6 kDa larger in molecular mass due to the myc and DDK epitopes with which these are tagged at their C-termini (Pin1-TRTRPL<u>EQKLISEEDL</u>AANDILDYK<u>DDDDK</u>V). Right panel: Quantification of PKCα steady-state levels in WT, Pin1 null cells, and Pin1 null cells reconstituted for expression of the indicated mutant Pin1 proteins as indicated at bottom. For quantification, actin was used to normalize the PKCα and Pin1 profiles for each cell line and the PKCα/actin ratios were set as 1.0 for parental WT Pin1 cells. Data represent the averages of five independent biological replicates ± standard error. Values were related to WT Pin1 controls using an unpaired two-tailed t-test (* p<0.05; **p<0.01; p***<0.001).

The CRISPR approach produced three HEK293T clones with little or no detectable Pin1 antigen, and those lines exhibited the highest steady-state levels of PKCα. Clone 3 represented a particularly attractive candidate for a Pin1 null cell line as it was devoid of detectable Pin1 antigen. This was confirmed by DNA sequencing. HEK293 cells carry four copies of the Pin1 gene (62), and DNA sequence analyses indicated clone 3 harbored three frameshift alleles. All three alleles altered the targeted exon 2 that encodes the Pin1 PPIase domain and included: (i) a 5bp deletion that interrupts the Pin1 amino acid sequence after Thr79, (ii) an 8bp insertion that interrupted the Pin1 sequence after Arg80, and (iii) a 10 bp deletion that disrupts the Pin1 amino acid sequence after Lys77 (**Fig. S19C**). The resulting translation products were prematurely terminated after addition of another twenty-two, ten and seven residues, respectively. Although PKCα steady-state levels were upregulated approximately 2-fold in clone 3 relative to parental wild-type HEK293T cells under both serum-free and serum-replete conditions (**Fig 7B**; 2.3 ± 0.2 and 2.3 ± 0.3-fold, respectively), the Pin1 null cells were not phenotypically perturbed. Their dimensions (diameter, surface area), viabilities and proliferation rates were indistinguishable from those of the parental cells (**Fig. S19D-G**). Thus, the elevation in steady-state PKCα levels identified a baseline Pin1 loss-of-function phenotype.

### Differential effects of Pin1 mutant expression on PKCα homeostasis *in vivo*

To assess the effects of Pin1 mutants on steady-state PKCα expression without the contribution of endogenous Pin1 activity, stable transgenic cell lines individually expressing epitope-tagged versions of WT Pin1; substrate-binding mutants: Pin1^W34A^, Pin1^R68A,R69A^, Pin1^W34A,R68A,R69A^; or the catalytic-deficient Pin1^C113S^ were derived from clone 3 cells. As shown in **Fig. 7C** (top left panel), WT Pin1 and all three of the Pin1 substrate binding mutants were over-expressed some 4-to 10-fold relative to endogenous Pin1 levels in these stable transgenic lines. Assessment of steady-state PKCα levels in the reconstituted cell lines showed that WT Pin1 expression significantly reduced PKCα steady-state levels relative to the Pin1 null condition (**Fig. 7C**, right panel). Reconstituted expression of Pin1^W34A^ exerted similar effects. By contrast, Pin1^R68A,R69A^ expression was less effective, and Pin1^W34A,R68A,R69A^ expression was completely ineffective, in restoring Pin1 function. As even significant enhancements (∼4-fold) in Pin1^W34A,R68A,R69A^ expression failed to downregulate PKCα levels (**Fig. 7C**, right panel), we conclude the triple mutant is strongly defective in PKCα regulation *in vivo*. Whereas Pin1^R68A,R69A^ scores as exhibiting greater functional deficiency than does Pin1^W34A^, more definitive interpretation of the Pin1^R68A,R69A^ and Pin1^W34A^ results is complicated by their overexpression. Elevated expression, particularly of Pin1^W34A^, likely compensates for partial binding defects to some degree and thereby obscures deficiencies in PKCα regulation *in vivo*.

Interestingly, we were unable to produce stable cell lines that overexpressed the Pin1^C113S^ ‘catalytic-dead’ mutant to the same levels achieved for WT Pin1 and the three substrate binding mutants. Reconstituted Pin1^C113S^ expression was consistently elevated only some 3-fold relative to endogenous Pin1 levels (**Fig. 7C**, bottom left panel). Yet, this relatively modest level of Pin1^C113S^ expression was at least as effective, if not more effective, than WT Pin1 expression in reducing steady-state PKCα levels in otherwise Pin1-null HEK293T cells (**Fig. 7C**, right panel). These collective results further support a physiologically relevant mechanism for Pin1-mediated regulation of PKCα that requires a bivalent interaction mode of Pin1 with PKCα, but operates independently of the Pin1 prolyl-isomerase catalytic activity.

## DISCUSSION

How Pin1 regulates the activities of its many *in vivo* substrates is of intense interest in contemporary biomedical science. This interest follows not only from the fact that the many targets of Pin1 regulation themselves play important cellular functions, but also that dysregulation of the Pin1 activity underlies the principal basis for multiple pathological disorders in humans. Pin1 has been the focus of many previous studies and it is generally accepted that prolyl *cis-trans* isomerase activity is its obligate functional feature. Yet, our understanding of how Pin1 engages its various substrates and regulates their activities remains incomplete. In this work, we provide a comprehensive biophysical, structural and cell biological description for how Pin1 recognizes/binds its PKCα and PKCβII substrates, and report the key features required for Pin1-mediated regulation of PKC degradation. Contrary to current dogma, we demonstrate a non-catalytic role for Pin1 in the regulation of PKC stability. We further show the underlying mechanism involves a bivalent interaction mode that has not been previously observed in any reported structures of Pin1 complexes. These discoveries not only expand the potential mechanisms by which Pin1 regulates the activities of its client substrates, but also hold interesting implications for the design of therapeutically effective Pin1 inhibitors.

### Proline at position +1 disqualifies Ser/Thr-Pro Pin1 binding motifs as isomerizable substrates

Previous experimental data suggested that Pin1-binding motifs with Pro at position +1 are disfavored substrates because of their failure to bind the catalytic PPIase domain of Pin1 (63). Our results provide direct experimental support for that conclusion, as both pTMα and pTMβII motifs bind with high affinities to the Pin1 WW domain but fail to bind the PPIase domain. In addition, ^1^H-^1^H exchange spectroscopy experiments clearly show that Pin1 is unable to catalyze the *cis*-*trans* isomerization of the pThr-Pro motifs of PKCα and PKCβII. The unsuitability of these motifs as Pin1 substrates is due solely to the presence of Pro at the +1 position as evidenced by our demonstrations that replacement this Pro residue restores their activities as substrates for isomerization by Pin1. Of note, the inability of Pin1 to catalyze *cis-trans* isomerization was previously observed for the pT^231^PP site in the Tau protein (52, 64). Interestingly, this observation prompted the speculation that Tau is not a genuine target for regulation by Pin1 – a speculation that our data indicate requires reconsideration (see below).

Why is Pro at position +1 incompatible with Pin1-catalyzed *cis-trans* isomerization of what is otherwise a signature substrate motif? Computational studies predict unfavorable overall binding free energies for Pin1 engagement with the transition state and the *cis*-conformations of the pTPP substrates (65). A definitive answer as to why Pro(+1)-containing motifs are incompatible with Pin1 catalysis remains elusive as there is currently no consensus on the precise catalytic mechanism of Pin1. The most recent ‘twisted-amide’ catalysis model (54) envisions a transition state where distortion of the substrate is stabilized by the H-bond between the C=O of the substrate Pro at position 0 and the backbone -NH of Pin1 residue Gln131. As formation and stabilization of the transition state relies on a dynamic network of H-bonds involving conserved residues in the Pin1 active site (59, 60, 66), Pro at position +1 might impose geometric constraints on the substrate where formation or stabilization of the amide twist is disfavored.

The demonstration that Pin1 S/TPP binding motifs are not isomerizable has interesting implications. A survey of AGC kinases reveals that these motifs are surprisingly common in this important class of enzymes. Moreover, as is the case for the PKC kinases, S/TPP motifs are present in the turn motif regions for both Akt and PKN kinases (**Fig. 8A**). These arrangements suggest that Pin1 binding to non-isomerizable substrates through its WW-domain defines an unappreciated, yet broadly deployed, regulatory strategy that operates independently of Pin1 *cis*-*trans* isomerase activity. This point is discussed in further detail below.

**Figure 8.**
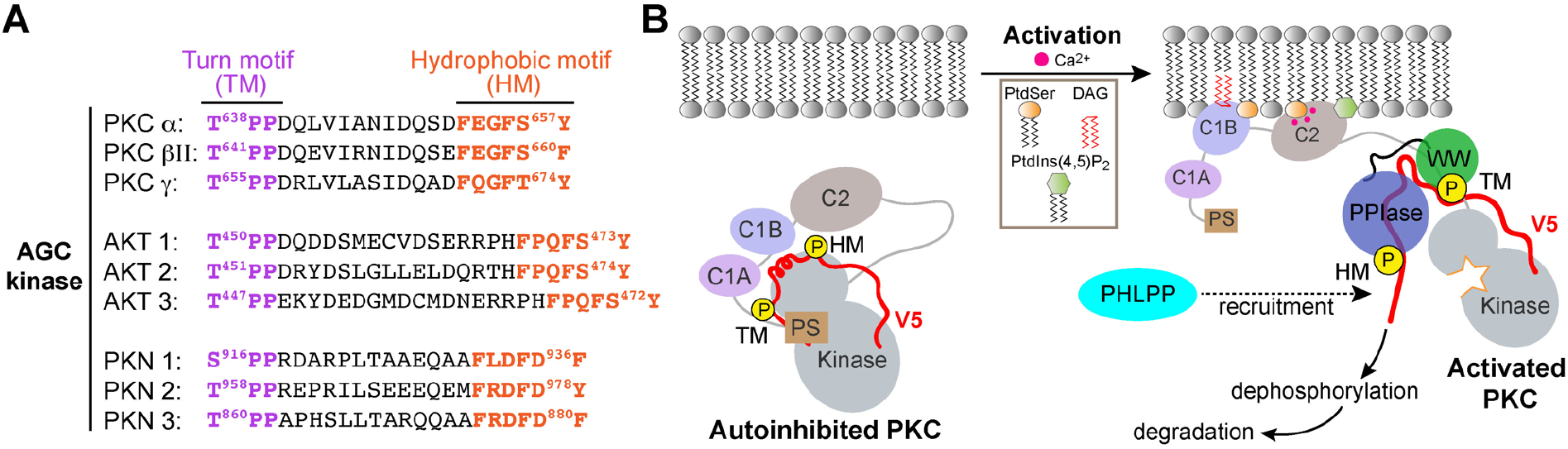
Non-catalytic role for Pin1. (**A**) Non-isomerizable pSer/Thr-Pro-Pro turn motifs separated by < 25 residues from the hydrophobic motifs are present in the C-term tails of other AGC kinases, such as AKT and PKN. (**B**) A possible model for Pin1-mediated downregulation of PKC α and βII isoforms. In the compact autoinhibited state, the pseudosubstrate (PS) blocks the catalytic site of the kinase and the C-terminal tail is not accessible. PKC activation involves Ca^2+^-dependent recruitment to the membranes where the regulatory domains, C1 through C2, bind diacylglycerol (DAG), phosphatidylserine (PtdSer), and phosphatidylinositol 4,5-bisphosphate (PtdIns(4,5)P_2_) and thereby trigger the release of autoinhibitory interactions. The activated open PKC conformation exposes the C-terminal V5 domain that is then engaged by Pin1 via bivalent binding to the phosphorylated turn motif (TM) and hydrophobic motif (HM). Pin1 might facilitate the recruitment of HM-specific PHLPP phosphatase to PKC by stabilizing the PKC open form. This promotes dephosphorylation of the hydrophobic motif by PHLPP and subsequent ubiquitination and degradation of the kinase.

### The Pin1-PKC interface is described by a novel bivalent interaction mode

Our NMR structure and NMR-detected binding experiments demonstrate that Pin1 interacts with the C-terminal tails of PKCα and PKCβII in a bivalent mode. The Pin1 WW-domain binds the TM, a canonical Pin1 pThr-Pro recognition motif, whereas the Pin1 PPIase-domain engages the HM, a non-canonical substrate that lacks Pro after the pSer residue. The specificity of TM binding exhibited by the WW-domain imposes a unidirectionality to the Pin1-PKC interaction by channeling the HM to the PPIase-domain. This bivalent binding mode results in Pin1 adopting a compact conformation, with the “pTM anchor” positioned at the interface between two Pin1 domains. Based on our structure, the linker length separating two phosphorylated Ser/Thr in bivalent Pin1 substrates is projected to be an important factor. In the PKCβII V5 domain, the phosphorylated TM and HM sites are separated by 18 amino acids. This length allows the simultaneous binding of the two sites by Pin1 without causing conformational strain on Pin1 itself. It is of interest to consider these results in light of what has been observed in the interaction of the yeast Pin1 homolog Ess1 with the RNA polymerase II C-terminal tail (67). The Ess1 linker that joins the WW- and PPase domains is a rather rigid element, and CTD repeats exceeding 27 amino acid residues are required to separate the two phosphorylated Ess1-binding sites for bivalent interactions (67). By contrast, the cognate Pin1 linker is unstructured, and its flexibility might afford Pin1 the potential to interact with a broader range of substrates.

Bivalent interactions afford significant advantages to biologically relevant protein-protein interactions as these are capable of increasing binding affinity and specificity, and inducing the appropriate conformational rearrangements in binding partners (68, 69). It is in this manner that the two substrate-binding domains of Pin1 govern its ability to engage in bivalent binding interactions that enhance the binding affinity and specificity to multi-phosphorylated client proteins (70, 71). Several Pin1 substrates that contain neighboring canonical pSer/Thr-Pro motifs in their unstructured regions have been characterized: IRAK1 kinase (71), Cdc25c (63), Tau (64), and STAT3 (72). Of those, the IRAK1-derived peptide was the only reported case of bivalent binding to Pin1 based on the NMR CSP data analysis, and this interaction involved two isomerizable pSer/Thr-Pro motifs (71). Our Pin1::pV5βII complex structure represents the first natural bivalent substrate-bound Pin1 structure. The two phosphate groups occupy the substrate binding site of the WW and PPIase domains, and the intervening residues stabilize the domain interface and bring the two domains into proximity. The linker that connects the Pin1 WW- and PPIase-domains maintains its flexibility and does not participate in interactions with the PKCβII C-terminal tail. The structure of this complex provides a guide for interpreting the potential binding modes of other Pin1 substrates that contain multiple pSer/Thr-Pro and non-canonical Pin1 motifs.

### Pin1 regulates PKC levels via a non-canonical non-catalytic mechanism

It is posited that Pin1 isomerase activity promotes ubiquitination and subsequent degradation of conventional PKC enzymes (23). This proposal rests on the observation that challenge of stimulated cells with a small molecule inhibitor of Pin1 abolishes agonist-induced ubiquitination of PKC. This conclusion is not ironclad, however, as it remains to be established that the inhibitor selectively targets isomerase activity without compromising the Pin1-PKC interaction. As described above, our collective biochemical and biophysical data suggest Pin1 downregulates PKC via a mechanism independent of its prolyl *cis*-*trans* isomerization activity. *In vivo* Pin1 reconstitution experiments provide direct support for this concept as expression of the ‘catalytic-dead’ Pin1^C113S^ mutant functionally rescues the Pin1 null condition and does so efficiently.

How might Pin1 operate via a non-catalytic mechanism to downregulate PKC activity? Upon activation, the membrane-associated conformation of PKC is sensitive to dephosphorylation – first at the HM site by <u>p</u>leckstrin <u>h</u>omology domain and leucine rich repeat <u>p</u>rotein <u>p</u>hosphatase (PHLPP) and subsequently at the TM site and activation loop by protein phosphatase 2A (PP2A) (21, 22, 73, 74). One possibility is that Pin1 discharges a scaffolding or substrate chaperoning function where it aids (directly or indirectly) in the recruitment of protein phosphatases to the Pin1-PKC complex. Such an activity, when coupled to a substrate chaperoning function, might ‘organize’ and stimulate ordered dephosphorylation of the PKC C-terminus with subsequent degradation of the kinase. Although Pin1 has not been demonstrated to interact with PHLPP or PP2A, such interactions are likely transient and difficult to capture. A recruitment mechanism of this nature is attractive in that it provides a means for channeling activated PKC to specific protein phosphatase(s) in a temporally and spatially appropriate manner (**Fig. 8B**). Alternatively, sequestration of the HM site by Pin1 might prevent the formation of the autoinhibitory state (75, 76) and thereby ‘trap’ PKC in an open “activated” conformation. This activated conformer would be particularly susceptible to the action of phosphatases that trigger its ultimate degradation. The significance of phosphorylated HM in maintaining the stability of AGC kinases is well documented (22, 74, 77).

It is tempting to speculate that non-catalytic functions for peptidyl-prolyl isomerases are more broadly represented in this group of enzymes. Pin1 and *E. coli* trigger factor (TF) are both peptidyl-prolyl *cis-trans* isomerases -- although Pin1 is a member of the Parvulin family of peptidyl-prolyl isomerases whereas TF is a member of the FK506-binding protein class. The enzymatic activity of TF is dispensable for its function as a protein folding chaperone (78). A TF^F198A^ mutant competent for client protein binding, but defective in peptidyl-prolyl *cis-trans* isomerization activity, remains active as a folding-promoting chaperone *in vivo* (78). Regardless, our data indicate one can no longer confidently infer whether a protein is a target of Pin1 regulation solely based on whether or not the putative substrate presents isomerizable pSer/Thr-Pro motifs.

### Implications for therapeutic interventions targeting Pin1

The oncogenic properties associated with dysregulation of Pin1 activities identify the isomerase as an attractive target for the development of new therapeutic approaches for cancer treatment (28, 79, 80). Unfortunately, although numerous small molecule Pin1 inhibitors have been identified (31, 81), these have been plagued by serious off-target toxicities. Resolution of this issue presents a significant obstacle to productive development of the Pin1-targeted drug pipeline (31, 82, 83). In that regard, the screens used to identify Pin1 inhibitors typically rely on readouts of catalytic activity. The data we report herein suggest alternative strategies. That is, screens for ligands that target the bivalent interaction mode rather than the Pin1 catalytic activity. Such strategies hold the potential for enhancing inhibitor specificity and affinity and thereby reducing the toxicities associated with off-target effects. The structure of the Pin1::pV5βII complex now offers a precise template for guiding the design and development of bivalent inhibitors that simultaneously bind to both the Pin1 WW- and PPIase-domains.

## METHODS

### 1. Protein expression and purification

A total of three Pin1 protein constructs were used in this study. The following genes with a codon-optimized DNA sequence were cloned into a pET-SUMO vector (Invitrogen): full-length Pin1 (residues 1-163), the WW domain (residues 1-50), and the PPIase domain (residues 50-163). Pin1 and isolated domains were heterologously expressed in *E.coli* BL21(DE3) and BL21(DE3)pLysS strains, respectively. For the natural abundance protein preparation, cells were grown in LB to an OD_600_ of 0.6 prior to induction of protein expression with 0.5 mM IPTG. The cells were grown for additional 4 to 5 hours at 37 °C. For the expression of isotopically enriched proteins, we used an LB to M9 minimal media re-suspension method (84). To generate uniformly ^15^N-enriched (U-[^15^N]) or ^15^N, ^13^C-enriched (U-[^15^N,^13^C]) protein samples, the M9 media contained either 1 g/L of ^15^NH_4_Cl and 3 g/L of natural-abundance D-glucose, or 1 g/L of ^15^NH_4_Cl and 3 g/L of ^13^C-D-glucose, respectively. The expression of isotopically enriched Pin1 in BL21(DE3) cells was induced for 15 hours at 15 °C. The expression of isotopically enriched individual domains in BL21(DE3) pLysS strain was induced for 5 hours at 37 °C.

The cells were harvested by centrifugation (4000 rpm, 30 minutes) at 4 °C. Cell pellets were resuspended in a buffer containing 20 mM Tris-HCl (pH 7.5), 0.5 M NaCl, 5 mM imidazole, and 10 mM β-mercaptoethanol. The 6×His-tagged SUMO fusion proteins were purified using a HisTrap™ HP Ni^2+^ affinity column (GE Healthcare Life Sciences). The fractions containing fusion protein were pooled and exchanged into a SUMO protease cleavage buffer (20 mM Tris-HCl at pH 8.0, 0.15 M NaCl) using a HiPrep 26/10 column (GE Healthcare Life Sciences). The cleavage reaction was initiated by adding 6×His-tagged SUMO protease to a final concentration of ∼8 μg/mL to the protein solution. After 30 minutes at room temperature, the reaction mixture was loaded onto to the second HisTrap™ HP Ni^2+^ affinity column to purify the desired protein from the 6×His-tagged SUMO and 6×His-tagged SUMO protease.

Purified proteins (Pin1, WW, or PPIase) were buffer-exchanged into an NMR buffer using either 5 or 3 kDa MWCO centrifugal concentrators (Vivaspin, Sartorius). The NMR buffer contained 10 mM d_4_-imidazole at pH 6.6, 100 mM KCl, 1 mM TCEP, 8% D_2_O, and 0.02% NaN_3_. The purity was assessed with SDS-PAGE conducted on samples with serial dilutions. Protein concentrations were determined by measuring the absorbance at 280 nm and using the following extinction coefficients: 20970 M^-1^cm^-1^ (Pin1), 13980 M^-1^cm^-1^ (WW), and 6990 M^-1^cm^-1^ (PPIase). The molecular masses of purified proteins were determined by MALDI-TOF mass spectrometry.

### 2. Synthesis and purification of the C-terminal PKC peptides

All 18 peptides used in this study were derived from the C-terminal V5 regions of either βII, α, or βI PKC isoenzymes (*H. sapiens*) and synthesized commercially (**Table S1**). The modifications included acetylation and amidation at the N- and C-termini, respectively, to avoid the influence of positive and negative charges terminal charges on binding and catalysis. All natural abundance peptides were purchased as crude mixtures and subsequently HPLC-purified on a C18 column (Waters). The buffers used for purification were 5 mM NH_4_HCO_3_ in 100% water (buffer A) and 5 mM NH_4_HCO_3_ in 82% acetonitrile/18% H_2_O (buffer B). A linear gradient of buffer B ranging from 0% to 20-40%, depending on the peptide, was applied during the elution step. The molecular weight and purity of the peptides were verified using ESI mass spectrometry. Isotopically enriched peptides, #14 and #15 in **Table S1**, were purchased in already purified form and used as is. For the binding experiments (*vide infra*), stock solutions of peptides were prepared by dissolving the lyophilized powders in ddH_2_O. The pH was adjusted to 6.6 either with NH_4_OH or HCl. The peptide concentration was determined using the absorbance at 205 nm; for the phosphorylated peptides, phosphate assay was additionally used (85).

### 3. Quantitative analysis of Pin1 interactions with the C-terminal PKC regions

NMR-detected binding experiments were conducted by adding aliquots of concentrated peptide stock solutions to the [U-^15^N] protein (Pin1, WW, or PPIase) samples, and recording [^15^N,^1^H] HSQC spectra for each peptide concentration point. The spectra were collected at 25 °C on Bruker Avance III spectrometers operating at the ^1^H Larmor frequencies of 600 or 500 MHz. The concentration of protein samples was between 90 and 100 µM. A total of 18 binding experiments were conducted; the parameters are summarized in **Table S2**. The binding data were analyzed in three ways, as described in sections A-C below.

#### 3A. Chemical shift analysis

The chemical shift perturbation analysis involved the calculation of the N-H_N_ cross-peak displacement between the two proteins sates, apo Pin1 and Pin1 in the presence of peptide at some specific concentration used in the binding experiments. The combined CSP value, Δ, was calculated in a residue-specific manner using the following equation:

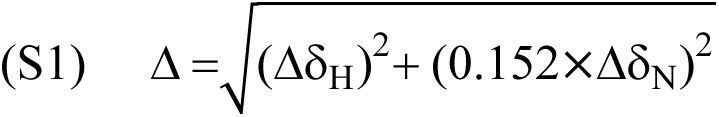

where Δδ_H_ and Δδ_N_ are the residue-specific differences in ^1^H_N_ and ^15^N chemical shifts, respectively.

#### 3B. Construction and analysis of the chemical shift-based binding curves

The binding curves for the affected residues were constructed by plotting Δ values against total peptide concentration. The cross-peaks with Δ values above the mean were used for quantitative analysis to extract the values of dissociation constant K_d_. Two binding models were used: single-site binding, and two independent sites binding.

##### Single-site binding model

In this model, the following equation was used for fitting the data is (86, 87):

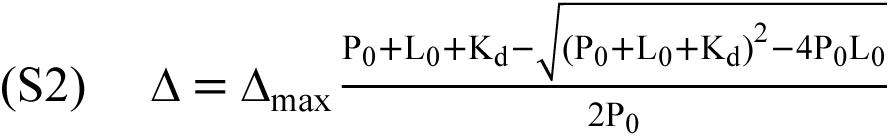

where K_d_ is the dissociation constant, Δ_max_ is the chemical shift changes at complete saturation, and P_0_ and L_0_ are total protein and ligand concentrations, respectively.

##### *2-site* binding model (independent sites)

According to our NMR data, the binding sites of the hydrophobic motif (pHM) on the Pin1 domains are independent, i.e. the binding to second site does not depend on whether or not first site is populated, and vice versa. The formalism outlined below is adapted from Wang et al (88). There are two binding equilibria that describe the process:

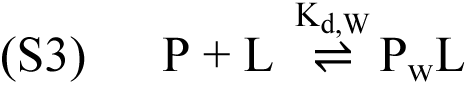

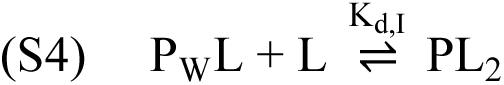

#### Definitions

P full-length Pin1

L ligand; in this case pHM

P_W_ Pin1 bound to L through the WW domain

P_I_ Pin1 bound to L through the PPIase domain

K_d,I_ dissociation constant for L binding to the PPIase domain K_d,W_ dissociation constant for L binding to the WW domain

A_W,i_ chemical shift difference between the apo and L-bound Pin1 for the i^th^ residue in the WW domain

A_I,i_ max chemical shift difference between the apo and L-bound Pin1 for the i^th^ residue in the PPIase domain

f_L,W_ fractional population of L-bound sites on the WW domain of Pin1 f_L,I_ fractional population of L-bound sites on the PPIase domain of Pin1

The chemical shift change due to ligand binding will be proportional to the fraction of the protein complexed to ligand L through a particular domain. In our case, we can equate this to the fractional population of ligand-bound sites of the WW and PPIase domains. These fractions can be expressed in terms of corresponding K_d_ values and <u>free ligand concentration:</u>

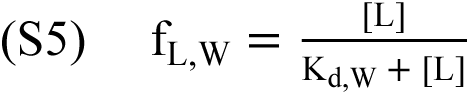

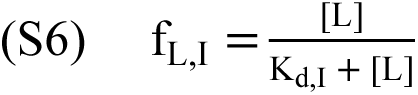

The expressions for the residue-specific chemical shift changes therefore become:

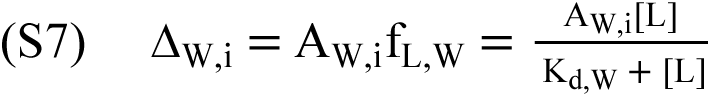

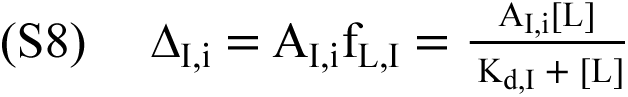

Wang et al. (88) provided the expression for [L] in terms of L_0_ and P_0_:

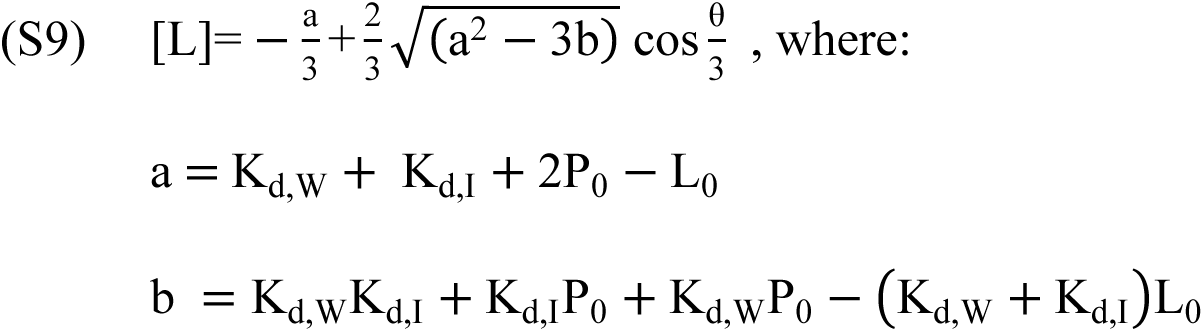

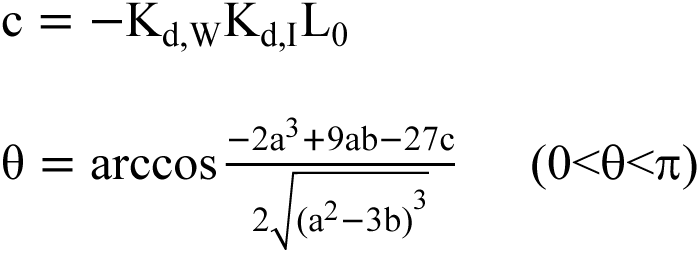

Equations (S7) and (S8), with the appropriate substitution of [L] from Eq. (S9), were used as fitting functions for the binding curves for the residues that belong to the WW and PPIase domains, respectively. To keep the fitting functions as simple as possible, we did not combine the ^1^H and ^15^N chemical shifts but used them individually to construct the experimental binding curves. The fitting was conducted using IgorPro software (Wavemetrics), with K_d,W_ and K_d,I_ as global parameters and Δ_W,i_ and Δ_I,i_ as the local ones.

#### 3C. Determination of binding affinities using lineshape analysis

K_d_ values that are determined from the chemical shift analysis become less accurate when the binding regime approaches the “tight” limit. Therefore, for the K_d_ values that are smaller that P_0_/5 (20 μM), we used the lineshape analysis to obtain the information about binding affinity. The lineshape analysis was conducted using the software package TITAN (89). For this analysis [^1^H, ^15^N]-HSQC were processed with exponential line broadening using NMRPipe (90). Resonances that are well resolved and are in fast exchange regime on the NMR chemical shift timescale were selected for the global fit using the “two-state ligand-binding” model, 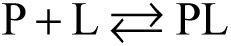, within TITAN. The input parameters for the analysis were the total protein concentration P_0_ (100 µM) and the total ligand concentration L_0_, the latter ranging from 0 to the saturating or near-saturating values. The parameters obtained from the fit were the dissociation constant K_d_ and the off-rate constant, k_off_.

### 4. ^1^H-^1^H exchange spectroscopy (EXSY)

The ability of Pin1 to catalyze the isomerization of the turn motif was probed using 2D ^1^H-^1^H exchange spectroscopy (EXSY) (91). The NMR samples contained 50 μM natural-abundance Pin1 and V5 peptides with concentrations ranging between 1 and 2 mM. The EXSY experiments were conducted on the total of six peptides: pV5βII, pV5α, pTMβII, pTMα, pTMα-P640A, and SP-2 (**Table S1**).

For the -pTPP-containing V5 regions, the maximum mixing time was set to 500 ms. The mixing times t_mix_ for pTMα-P640A were: 20, 35, 50, 75, 100, 140, 200, 280, 400, and 500 ms. The mixing times t_mix_ for the SP-2 were: 10, 25, 30, 40, 70, 100, 150, 200, 300, 400, and 500 ms. The *cis*-*trans* isomerization of the pThr-Pro bond was monitored by following the intensity of the diagonal- and cross-peaks corresponding to the ^1^H_N_ resonance of Thr641 and Thr638 in the βII and α peptides, respectively. The time-dependence of the ratio of intensities of the cis→trans (I_ct_) and diagonal trans-trans peak (I_tt_), I_ct_/I_tt_, is given by the following equation:

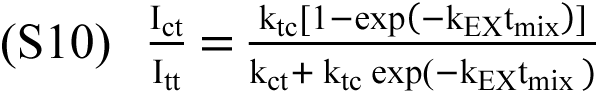

where k_ct_ and k_tc_ are the forward and reverse rate constants for cis→trans isomerization reaction, k_EX_ is the exchange rate constant that is equal to k_ct_+k_tc_, and t_mix_ is the mixing time. Eq. (S10) was used to fit the data, with k_tc_ and k_ct_ as the adjustable parameters. The populations of the cis- and trans-conformers of the peptidyl-prolyl pThr-Pro bond were estimated using the SP-2 peptide (LpTPTD), where the cis-cis diagonal peak is well resolved. The populations of the trans- and cis-conformers of the pThr-Pro bond were I_tt_/(I_tt_+I_cc_) = 91% and I_cc_/(I_tt_+I_cc_) = 9%, respectively.

### 5. Determination of the Pin1-pV5βII complex structure using solution NMR spectroscopy 5A. Sample preparation

We prepared a total of 10 NMR samples for the structural analysis of the Pin1-pV5βII complex. The samples are listed in **Table S3**, along with the corresponding NMR experiments. The NMR buffer was 10 mM ^2^H_4_-imidazole at pH 6.6, 100 mM KCl, 1mM TCEP, 8% D_2_O, and 0.02% NaN_3_, except for sample #2* where 100% D_2_O was used. Samples #1-3 were used to collect the NMR data used for resonance assignments and structure determination. Samples #4-10 provided validation for the assignments. The validation relied on the spectral simplification through the use of single-domain constructs of Pin1 and selectively labeled pTM and pHM regions of pV5βII. All 2D and 3D NMR experiments were acquired at 25 °C on the Bruker Avance III HD spectrometers operating at the ^1^H Larmor frequency of 600 or 800 MHz. The temperature was calibrated using deuterated methanol. The chemical shifts were externally referenced to DSS. The NMR data were processed with NMRPipe (90) and analyzed with CcpNmr-Analysis program, version 2.4.2 (92).

#### 5B. NMR structure calculation and refinement

The backbone and sidechain ^1^H resonances of Pin1 complexed to pV5βII were assigned using standard 3D NMR methods (**Table S3**). Pin1-bound pV5βII ^1^H resonances were assigned using 2D-double filtered NOESY and TOCSY experiments conducted on samples #3 and #5 (**Table S3**). To facilitate the assignments and overcome the problem of extensive spectral overlap, we prepared NMR samples where the pTMβII and pHMβII peptides were selectively labeled with [U-^13^C,^15^N] amino acids at specific positions (**Table S1**, peptides #14 and #15; **Table S3**, samples #6 and #7).

The distance restraints were obtained from the height of the ^1^H-^1^H cross peaks in the NOESY spectra. The inter-molecular NOEs were obtained using 3D-^15^N/^13^C-edited ^13^C, ^15^N-filtered NOESY-HSQC (93, 94) on samples #1, #2, #6 and #7; and 2D [F1/F2] ^13^C, ^15^N-filtered NOESY on sample #3. A total of 2812 (662 of them long-range) intra-Pin1, 241 intra-pV5βII, and 75 inter-molecular Pin1-pV5βII NOEs were used for the structure calculation (**Table S4**). Hydrogen bonds were identified based on the ^1^H-^2^D exchange rates of amide ^1^H atoms. Dihedral angles were predicted by the DANGLE routine within the CcpNmr program (92) using a complete set of ^15^N, ^13^C′, ^13^C^α^, ^13^C^β^, ^1^H^α^ and ^1^H^N^ chemical shifts.

An ensemble of the Pin1-pV5βII complex structures was calculated using the torsion angle dynamics protocol in CYANA (version 3) (95, 96). Nine hundred random conformers were subjected to 20000 steps of annealing. The fifty low-energy conformers that had no upper distance violations of >0.2 Å or dihedral angle violations of >5° were used as an input for the refinement procedure implemented in Xplor NIH, version 2.51.5 (97, 98). The refinement was conducted in explicit solvent, using the TIP3P water model. The refined ensemble comprising 20 structures was deposited in the PDB under the accession code 8SG2.

### 6. In-cell experiments 6A. Reagents

Chemical reagents were obtained from MilliporeSigma (Burlington MA) or Thermo Fisher Scientific (Waltham, MA) unless otherwise specified. Disposable plastics, tissue culture dishes, etc. were from Genesee (El Cajon, CA) or VWR (Radnor, PA). Primary antibodies obtained from Cell Signaling Technology® (Denvers, MA) included those directed against Pin1 [3722S], beta actin [4970S], PKCα [20565], PKCβ [46809S], and GFP [2555S]. Primary anti-DDK immunoglobulin [TA50011-100] was obtained from OriGene Technologies Inc. (Rockville, MD). Secondary goat anti-rabbit IgG (H+L) HRP conjugated antibody was from MilliporeSigma. HEK293T cells were purchased from the American Type Culture Collection (CRL-3216™, ATCC, Manassas, Virginia, USA).

#### 6B. Generation of Pin1 mutant expression plasmids

A Myc/DDK eptiope-tagged Pin1 (NM_006221) Human Tagged ORF Clone (Cat# RC202543) was obtained from OriGene Technologies Inc. The Q5^®^ Site-Directed Mutagenesis Kit (NEB, Ipswich, MA) was used to generate the appropriate Pin1 point mutants: Pin1 W34A, R68A/ R69A, C113S, and W34A/R68A/ R69A. All site-directed mutations were verified by DNA sequence analysis.

The primer sequences for the construction of each point mutant are as follows (sites of mutation are highlighted):

W34A:    forward primer 5’- CGCCAGCCAG**G**CCGAGCGGCCCA −3’

          reverse primer 5’- TTAGTGATGTGGTTGAAGTAGTACAC −3’

R68A/R69A:     forward primer 5’- CAGCCAGTCA**GCCGCC**CCCTCGTCCTGGCG −3’

          reverse primer 5’- TGCTTCACCAGCAGGTGC −3’

C113S:     forward primer 5’- GTTCAGCGAC**A**GCAGCTCAGCCA −3’

          reverse primer 5’- TGTGAGGCCAGAGACTCAAAG −3’

#### 6C. Cell culture, and plasmid transfections

HEK293T cells were cultured in a humified incubator at 37°C and 5% CO_2_ in high-glucose Dulbecco’s Modified Eagle Medium (DMEM) plus sodium pyruvate (Genesee) and supplemented with 10% FBS and penicillin (100 units/ml)/streptomycin (100µg/ml). After trypsinization to release adherent cells, viable cell counts and cell size were determined by trypan blue staining (Invitrogen, Thermo Fisher Scientific) followed by passage through a Countess II^TM^ automated cell counter (Thermo Fisher Scientific) according to manufacturer’s instructions. Plasmid transfections were performed using Lipofectamine™ LTX with Plus reagent™ (Thermo Fisher Scientific) according to manufacturer’s instructions. Stable expression lines were generated by transfection followed by serial selection with 2mg/ml geneticin (Gibco).

#### 6D. Cell lysate preparation and Pin1 immunoblotting

Expression of Pin1, its corresponding variants, and PKCα levels were monitored by immunoblotting. Cells were solubilized in lysis buffer (BB150: 50 mM Tris pH 7.6, 150 mM NaCl, 0.2% CHAPS, 10 mM EDTA) on ice and centrifuged at 21,000xg for 5min at 4°C to remove insoluble materials. Protein in the soluble fractions was quantified using the colorimetric bicinchoninic acid assay (Pierce) and samples were prepared with 80µg of total lysate protein in 1x Laemlli buffer (62.5 mM Tris-HCl, 2% SDS, 25% glycerol, and 0.01% bromophenol blue, pH 6.8). Samples were subsequently boiled for 5min and resolved by SDS-PAGE using 15% acrylamide gels (120V constant voltage for 80 min) and transferred onto 0.2 µm nitrocellulose membranes (Bio Rad) at 350 mA (constant current) for 1h. Following transfer, membranes were blocked for 1h in 5% milk powder (w/v) in 20 mM Tris pH7.6, 150 mM NaCl, 0.1% Tween 20 (TBST), and incubated at 4°C overnight with primary antibodies at a 1:1000 dilution in Tris-buffered saline (pH7.6), 0.1% Tween 20, 5% BSA (w/v) with rocking. Following incubation, blots were washed three times for 10 min each in TBST and incubated with goat anti-rabbit secondary antibody coupled to horseradish peroxidase (MillliporeSigma; 1:10,000 dilution) in TBST/5% BSA for 1h at room temperature. Following three additional 10 min washes with TBST, blots were developed using SuperSignal™ West Femto Maximum Sensitivity Substrate (Thermo Fisher Scientific) and analyzed by densitometry using a Molecular Imager® ChemiDoc™ XRS+ system (Bio-Rad Laboratories, Hercules, CA). Protein levels were normalized to total protein load and verified using actin as loading control.

#### 6D. Quantification of PKCα levels

PKCα levels were assessed in cells seeded at 1.5 x10^6^ cells per 35 mm dish 24h prior to analysis. Cells from three dishes were pooled per sample. Cells were seeded in complete media containing 10% FBS unless otherwise stated. Following incubation, cells were washed with phosphate buffered saline and snap frozen in liquid nitrogen prior to analysis.

#### 6E. siRNA-Mediated knockdown of Pin1

siRNA transfections were performed using Dharmafect (Dharmacon/ Horizon) according to manufacturer’s instructions in antibiotic free growth media. HEK293T cells were incubated with siRNA complexes overnight and complete media were exchanged the following day. Optimal Pin1 knockdown was determined to be at 72h post-transfection. All transfection complexes for both sets of reagents were prepared in optimum media lacking FBS or antibiotics. The ON-TARGET plus Human Pin1 (5300) siRNA-SMART pool (Dharmacon/ Horizon) sequences were as follows: J-003291-10 5’-GCUCAGGCCGAGUGUACUA−3’, J-003291-11 5’-GAAGACGCCU CGUUUGCGC−3’, J-003291-12 5’-GAAGAUCACCCGGACCAAG−3’, and J-003291-13 5’-CCAC AUCACUAACGCCAGC−3’.

#### 6F. Isolation of a clonally-derived Pin1 null cell line

To generate clonal cell lines lacking Pin1 protein, HEK293T cells were sequentially transfected with two CRISPR/Cas9 plasmids driving expression of guide RNAs that specifically target exon2 of the *Pin1* structural gene (MilliporeSigma product numbers and targeting sequences: CRISPRD HSPD0000031439 5’-GA GAAGATCACCCGGACCA−3’, CRISPRD HSPD0000031440 5’-TAACGCCAGCCAGTGGG AG−3’). Transfected cells were flow-sorted on the basis of GFP fluorescence on a 3-laser (405nm, 488nm, 640nm) Beckman Coulter Moflo Astrios high-speed cell sorter at a rate of < 1000 events/sec. Cell debris were gated and eliminated from the bulk sort. GFP was detected using a 664/22 nm bandpass filter, and Cas9-GFP positive cells were bulk-sorted into 1.5ml tube using the ‘Purify mode’ routine with a drop envelope of 1-2 droplets.

Following an initial round of sorting, cells were allowed to recover, and the bulk-sorted population was re-transfected with the second CRISPR/Cas9 plasmid and re-sorted as described. Those second-round bulk-sorted cells were tested to assess overall Pin1 expression within the heterogeneous cell population by immunoblotting for Pin1 antigen. In procedures where the bulk population expressed ≤ 50% of wild-type Pin1 levels using total protein to normalize data, individual cell lines were developed from the bulk-sort population by limiting serial dilution and expansion of single cells into clonal cell populations. Clonally-derived cell lines lacking detectable Pin1 antigen were identified by immunoblotting with anti-Pin1 immunoglobulin.

Clonally-derived cell lines were verified by sequencing of individual *Pin1* alleles. Total genomic DNA was isolated from each cell line using the GenElute™ Mammalian Genomic Miniprep Kits (MilliporeSigma). The *Pin1* coding region from exons 2 and 3 was recovered by amplifying each of these coding exons individually via polymerase chain reaction using the Phusion-Plus high-fidelity proof-reading enzyme (ThermoFisher Scientific). Primers used for amplification of each exon were as follows:

Exon 2 -- Forward 5’-CTGGGAGCACAACCCTAGC−3’ Reverse 5’-AGGTCATGCACTGGCGTTTT−3 Forward 5’-GGGAGCACAACCCTAGCTG−3 Reverse 5’-CTACAAAGGCTCACCTGGGA−3

Exon 3 -- Forward 5’-CTGGCACTCCCATTCCGTTC−3 Reverse 5’-CCTGCCATGTCATCTGTCCC−3 Forward 5’-ACTCCCATTCCGTTCCATGTC−3 Reverse 5’-CCCTGCCATGTCATCTGTCC−3

Exon 4 -- Forward 5’-CAGGTCAGATGCAGAAGCCAT−3 Reverse 5’-CCACGACATCTTCCCCACTAT−3 Forward 5’-AGGTCAGATGCAGAAGCCATT−3 Reverse 5’-GATCCCCTCCCCACGACATC−3

PCR products were extended by *Taq* polymerase-driven poly-deoxyadenosine tailing in 30 min. Tailed PCR products were analyzed by agarose gel electrophoresis alongside products generated from a wild type HEK293T cell control. Bands were excised and gel purified using a Qiaquick® Gel Extraction Kit (Qiagen). Purified DNA fragments were ligated into the pGEM®t-easy plasmid (Promega) and transformed into DH5α bacteria selected on standard Luria broth (LB) agar plates containing ampicillin (100 µg/ml), X-gal (50 µg/ml) and IPTG (1 mM). Blue/white colony screening was employed to identify those transformants harboring plasmids containing inserts (white colonies). Plasmids were prepared from individual colonies using the QIAprep® Spin Miniprep Kit and individual allele sequences were obtained by DNA sequencing using the T7 primer to program the sequencing reaction.

#### 6G. Production of stable transgenic cell lines expressing Pin1 variants

To generate cell lines expressing wild-type or mutant Pin1 proteins as sole sources for Pin1 activity, HEK293T Pin1 KO cells (clone C3) were transfected with WT Pin1, the W34A, R68A/R69A, W34A/R68A/R69A, or C113S transgenes tagged with the myc/DDK epitopes. The epitope tags were engineered at the Pin1 C-terminus using the following DNA sequence (myc and DDK coding sequences underlined):

5’- ACG CGT ACG CGG CCG CTC GAG CAG AAA CTC ATC TCA GAA GAG GAT CTG GCA GCA AAT GAT ATC CTG GAT TAC AAG GAT GAC GAC GAT AAG GTT TAA −3’

Twenty-four hours post-transfection, an aliquot of the transfected cells was harvested to generate lysates from which successful transfection and protein expression were assessed by immunoblotting for Pin1 antigen. The remaining cells were cultured in high-selection media (2 mg/ml geneticin) and grown at low density, with repeated passage and frequent media changes, over a period of three weeks. Pin1 expression was subsequently re-examined by immunoblotting for Pin1 antigen and verified stable lines were consistently maintained under low-selection conditions (1 mg/ml geneticin).

#### 6H. Cell area and cell viability measurements by flow cytometry

Cells were loaded into the Image Stream X Mark II imaging flow cytometer (Cytek/Amnis Freemont, CA) to measure cell area and viability. Cells were analyzed at a concentration of 1×10^6^ cells in 50 µl of FACS buffer (phosphate buffered saline, 0.5% bovine serum albumin) supplemented with eBioscience™ propridium iodide (Invitrogen) to a final concentration of 2µg/ml. All data from the Image Stream were acquired using the Inspire program (version 201.1.0.765; Cytek/Amnis Freemont, CA). The acquisition parameters used were 40x objective, medium speed, 488nm wavelength, laser power 100mW, SSC laser power 2.81 mW. The scatter channel was set for channel 6, the bright field channels were set to channels 1 and 9. The core stream size was set to 6 µm, and 10,000 events of focused, live, single cells were collected. The focus was determined by plotting the Gradient RMS values for channel 1 bright field in a histogram and selecting cells that possessed a gradient RMS value of ≥ 40. Single cells were identified in the focused cells by plotting the aspect ratio of the channel 1 bright field (y-axis) versus channel 1 bright field area (x-axis). Single cells reside in the range of 0.6 to 1 in the aspect ratio and between 0 and 1×10^3^ µm^2^ range in the area of channel 1. Cell viability was determined by plotting the intensity of channel 5 (propidium iodide) on the y-axis versus intensity of channel six (side scatter). The data were analyzed using the acquisition gates and settings in the IDEAS 6.3 (Cytek/Amnis Freemont, CA) data analysis software package. The median of the cell area and % viability parameters were calculated for each sample.

## AUTHOR CONTRIBUTIONS

TII and YY conceived the project. XRC, KD, YY, and HTI prepared the samples, conducted NMR experiments, and analyzed the data. KD determined the structure of the Pin1-C-term PKCβII complex. XRC and TII conducted the structural analysis. MIM and VAB designed *in vivo* experiments. MIM performed *in vivo* experiments and MIM and VAB analyzed the data. All authors contributed to the preparation of the manuscript.

## ACKNOWLEDGEMENTS

This work was supported by grants NIH RO1 GM108998 and Robert A. Welch Foundation A-1784 to TII. YY was an American Heart Association predoctoral fellow (award no. 14PRE20380475). MIM and VAB were supported by grants NIH R35 GM131804 and award BE-0017 from the Robert A. Welch Foundation to VAB. We acknowledge Gus Wright (Texas A&M Veterinary School Flow Cytometry Core) for assistance in the cell sorting component of the PIN1 gene editing experiments and with cell viability and cell area measurements.

## DATA AVAILABILITY

The structural data of coordinates, restraints, chemical shifts, and peak lists for the Pin1::pV5βII complex are deposited in the wwPDB under the accession codes: 8SG2 and BMRB-31080.

