## Supporting information for "A novel bivalent interaction mode underlies a non-catalytic mechanism for Pin1-mediated Protein Kinase C regulation"

### FIGURES

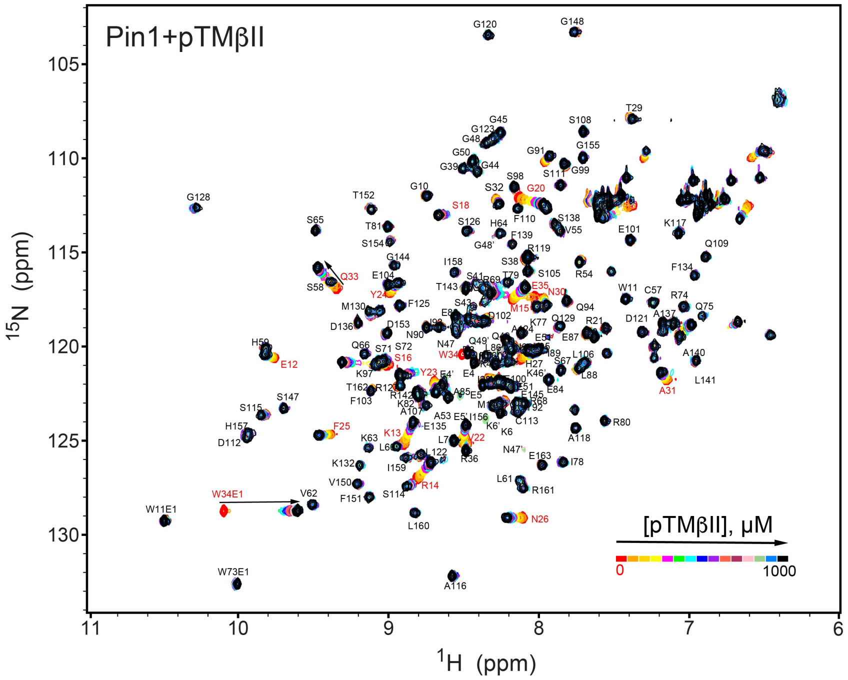

**Figure S1. NMR-detected binding of the PKCβII turn motif (pTMβII) to full-length Pin1.** The protein concentration was 100 μM, and the pTMβII concentration varied from 0 to 1 mM. Most of the affected N-$H_N$ resonances are in fast exchange on the NMR chemical shift timescale (**Table S2,** binding experiment #1).

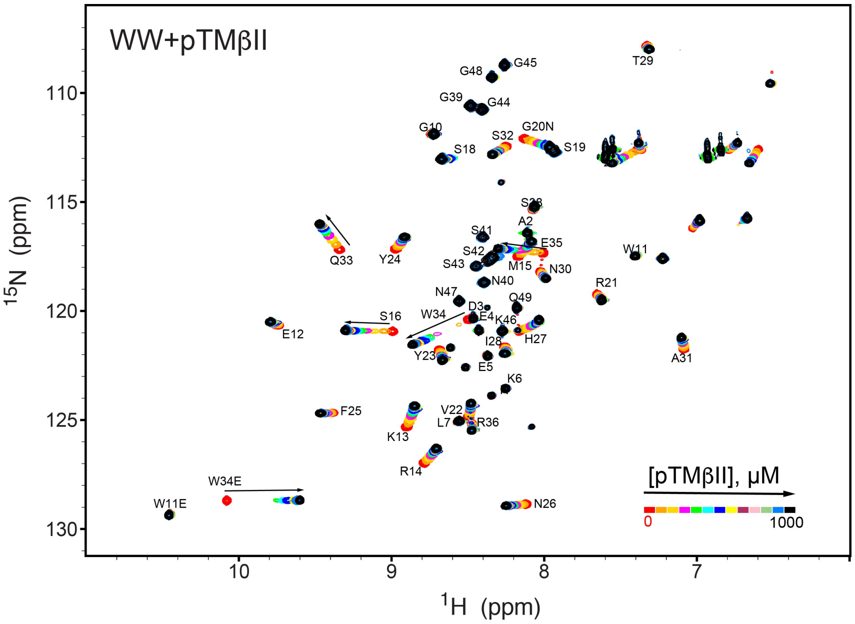

**Figure S2. NMR-detected binding of the PKCβII turn motif (pTMβII) to the isolated WW domain.** The protein concentration was 100 μM, and the pTMβII concentration varied from 0 to 1 mM. Most of the affected N-$H_N$ resonances are in fast exchange on the NMR chemical shift timescale (**Table S2,** binding experiment #2).

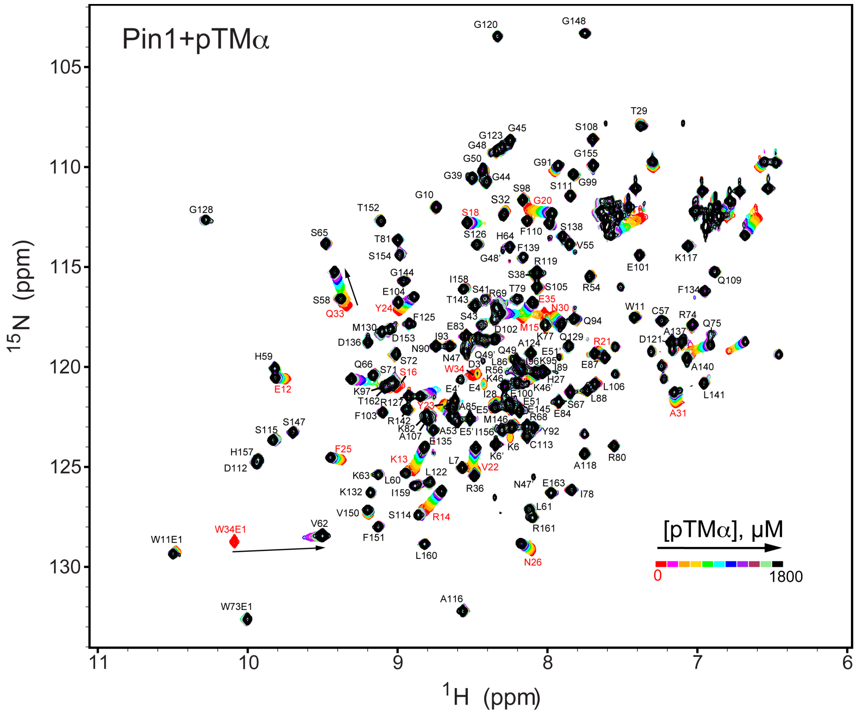

**Figure S3. NMR-detected binding of the PKCα turn motif (pTMα) to full-length Pin1.** The protein concentration was 100 μM, and the pTMα concentration varied from 0 to 1.8 mM. Most of the affected N-$H_N$ resonances are in fast exchange on the NMR chemical shift timescale (**Table S2,** binding experiment #3).

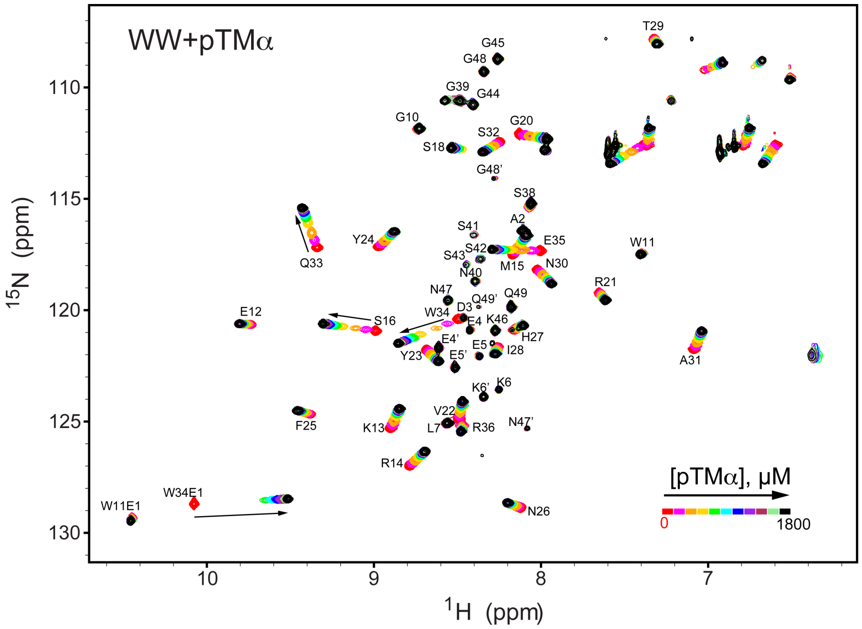

**Figure S4. NMR-detected binding of the PKCα turn motif (pTMα) to the isolated WW domain.** The protein concentration was 100 μM, and the pTMα concentration varied from 0 to 1.8 mM. Most of the affected N-$H_N$ resonances are in fast exchange on the NMR chemical shift timescale (**Table S2,** binding experiment #4).

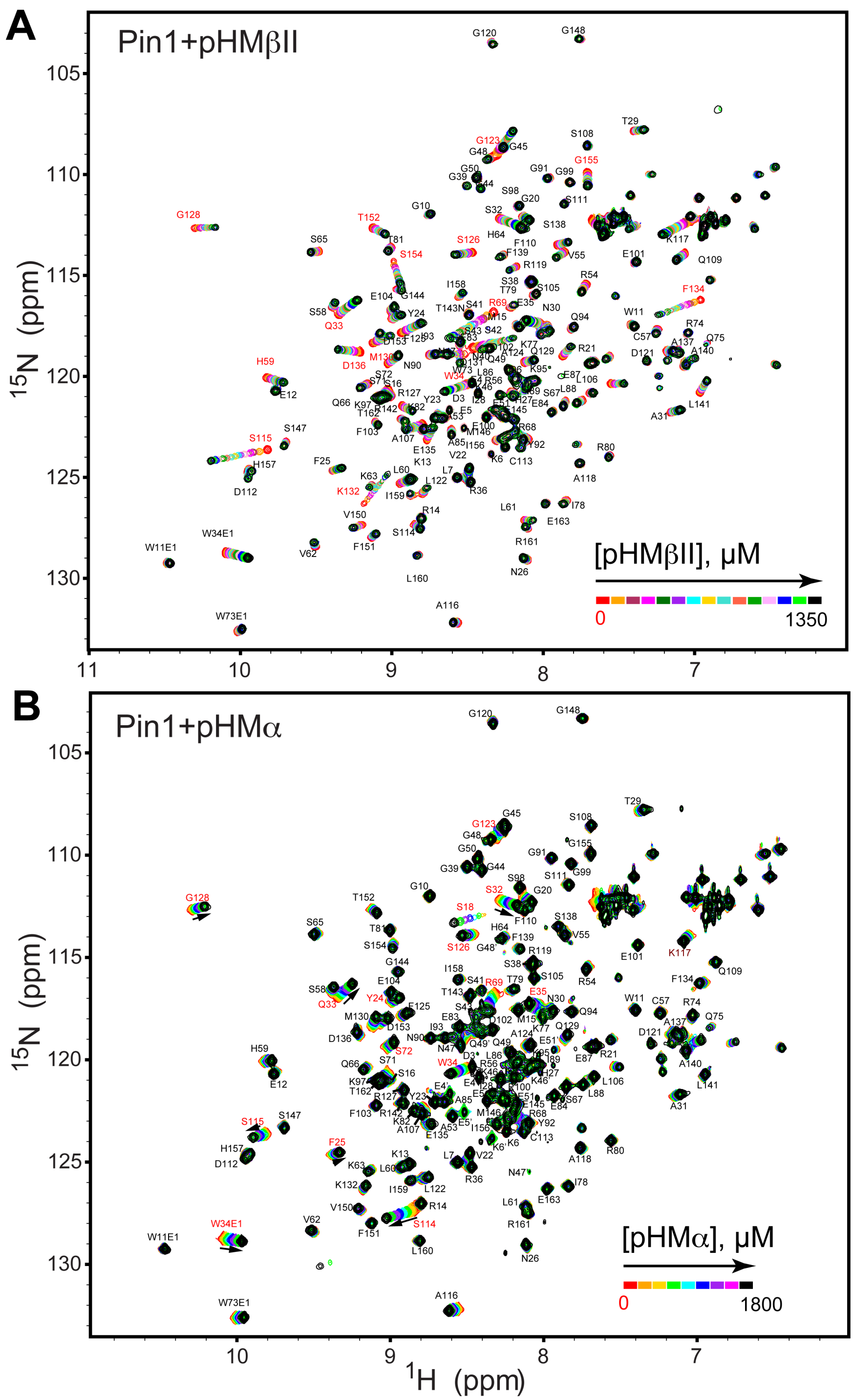

**Figure S5. NMR-detected binding of hydrophobic motifs from PKCβII (A, pHMβII) and α (B, pHMα) to full-length Pin1.** The protein concentration was 100 μM, and the pHM concentration varied from 0 to 1.35 mM (pHMβII), and from 0 to 1.8 mM (pHMα). Most of the affected N-$H_N$ resonances are in fast exchange on the NMR chemical shift timescale (**Table S2,** binding experiments #7 and #10).

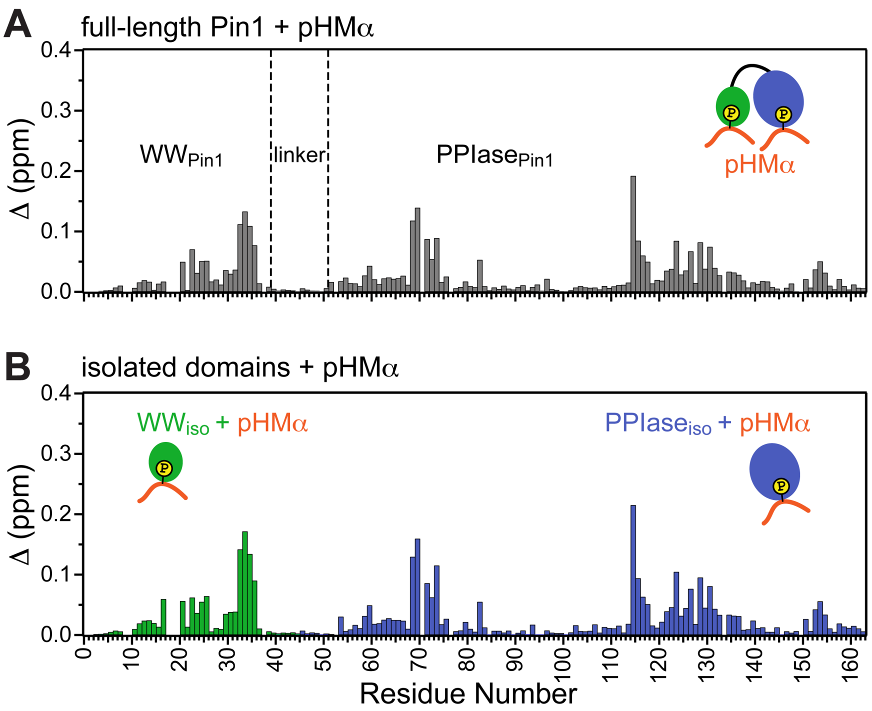

**Figure S6. Pin1 binds hydrophobic motif of PKCα (pHMα) via two independent sites.** Comparison of the CSP plots obtained at maximum concentrations of pHMα used in binding experiments versus ligand-free proteins for (**A**) full-length Pin1 and (**B**) isolated WW and PPIase domains. The similarity of the CSP patterns in (**A**) and (**B**) indicates that Pin1 has two pHM binding sites, one per domain. The protein concentration was 100 μM. Other details are given in **Table S2** for the binding experiments #5, 6, and 7.

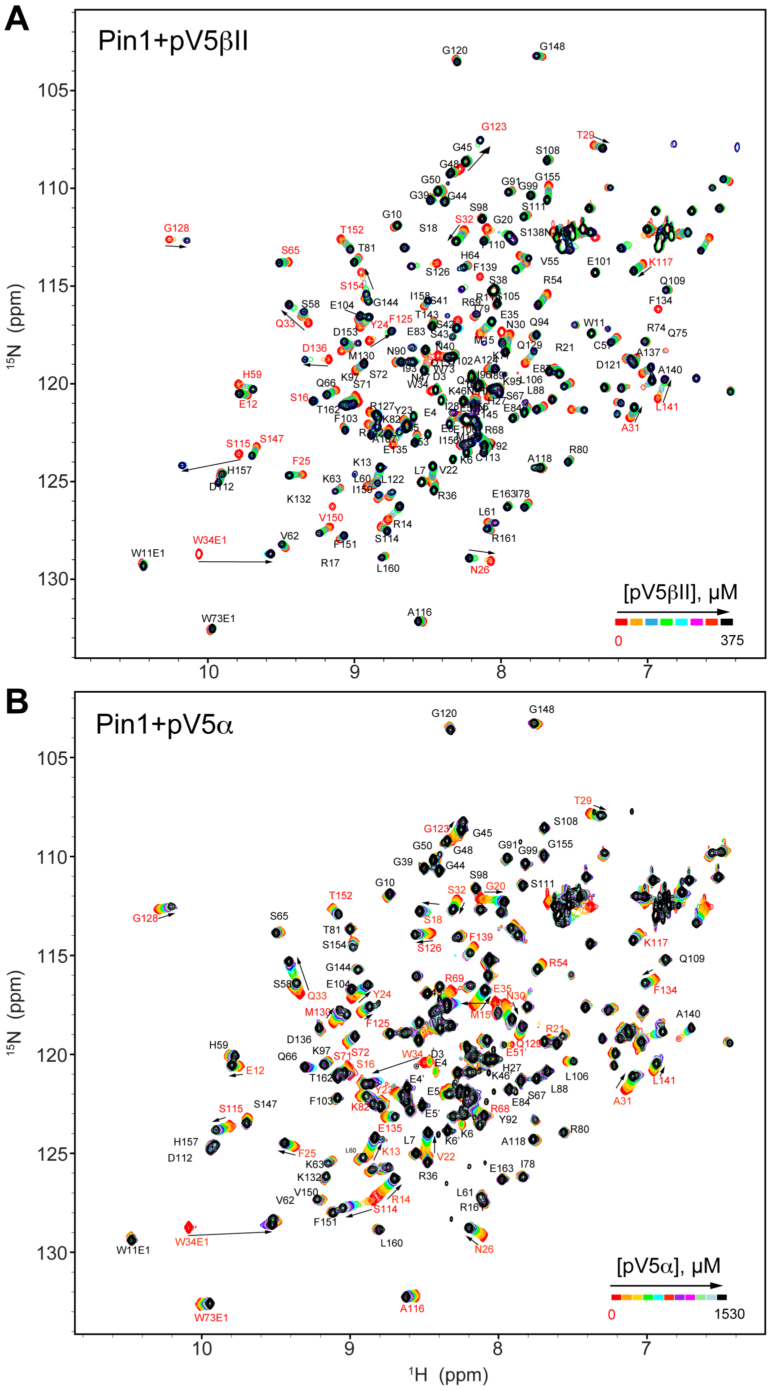

**Figure S7. NMR-detected binding of the fully phosphorylated C-term regions pV5βII (A) and pV5α (B) to full-length Pin1.** The protein concentration was 100 μM, and the pV5 concentration varied from 0 to 375 μM in case of pV5βII, and from 0 to 1530 μM in case of pV5α. The binding kinetics is in the intermediate-to-fast (pV5βII) and fast (pV5α) exchange regimes (**Table S2,** binding experiments #14 and #15).

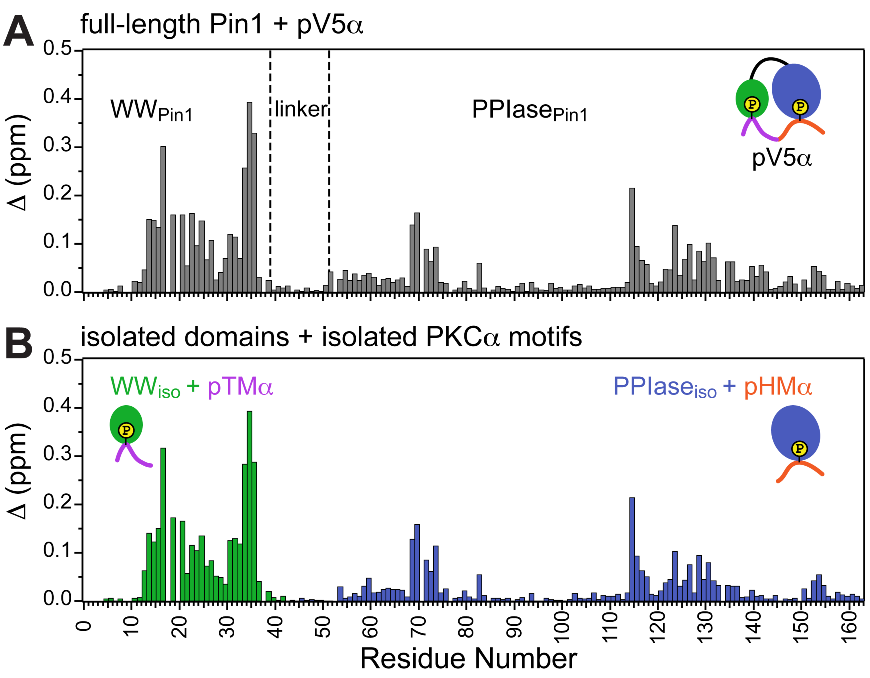

**Figure S8. Unidirectional bivalent binding mode of the C-terminal PKCα region to Pin1.** Comparison of the CSP plots of Pin1 obtained at maximum concentrations of pV5α (**A**) and those of isolated domains, $WW_{iso}$ and $PPIase_{iso}$, at maximum concentrations of pTMα and pHMα (**B**), respectively. The similarity of CSP patterns in (**A**) and (**B**) indicates that the C-term region of PKCα binds to Pin1 in a unidirectional bivalent mode. The TM and HM binding sites reside on the WW and PPIase domains, respectively. The protein concentration was 100 μM. Other details are given in **Table S2**, binding experiments #4, 6, and 14.

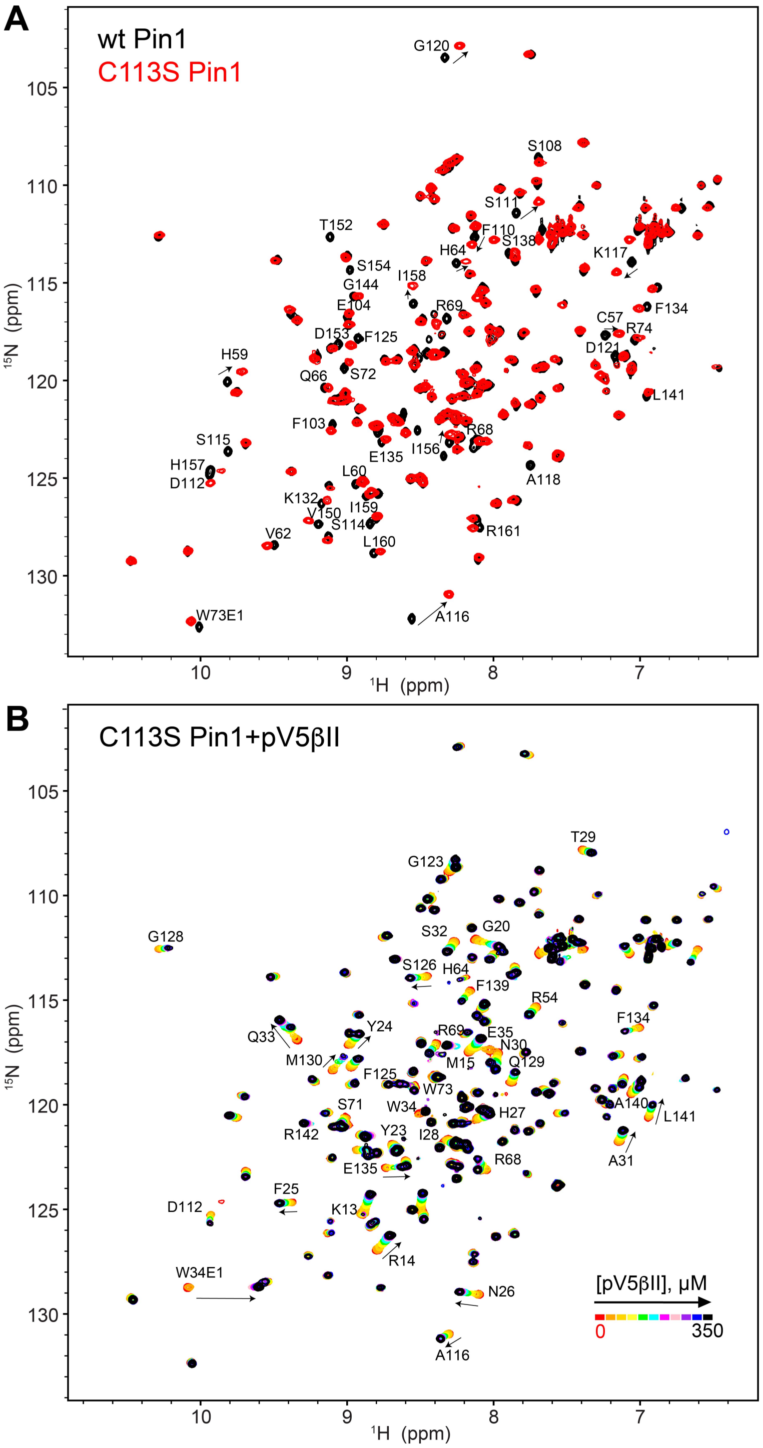

**Figure S9. The C-term region of PKCβII binds to the catalytically deficient C113S Pin1 variant. (A)** The C113S Pin1 spectrum (red) shows minimum chemical shift perturbations compared to that of the wild-type Pin1 (black). **(B)** The C-term region pV5βII binds to C113S Pin1, evidenced by the chemical shift changes upon addition of increasing amounts of pV5βII. The chemical exchange regime is identical to that observed for the wt Pin1 in **Fig. S7A**. The protein concentration was 100 μM, and the pV5βII concentration varies from 0 to 350 μM (**Table S2**, binding experiment ID #19).

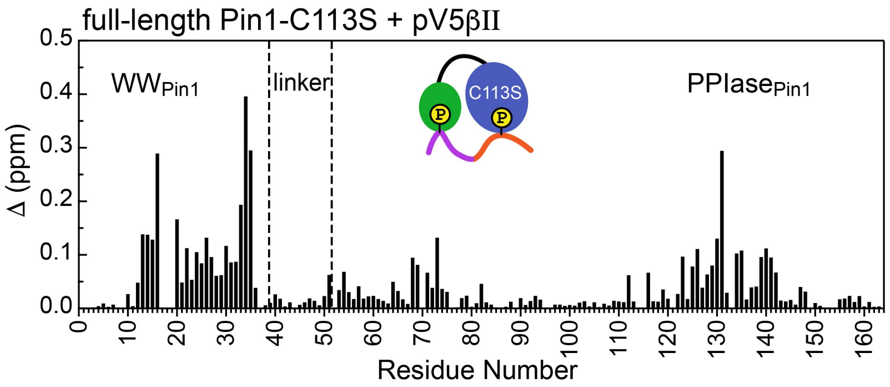

**Figure S10. Unidirectional bivalent binding mode of the C-terminal PKCβII region to C113S Pin1.** The CSP plot was constructed using the chemical shifts of the apo and pV5βII-bound C113S Pin1. The similarity of CSP patterns between the pV5βII-complexed wild-type (**Fig. 4A** of the main manuscript) and C113S Pin1 indicates that the binding mode of the PKCβII C-term region does not change as a result of the mutation. The protein concentration in the C113S Pin1 experiments was 100 μM and the maximum concentration of pV5βII was 350 μM (**Table S2,** binding experiment #19).

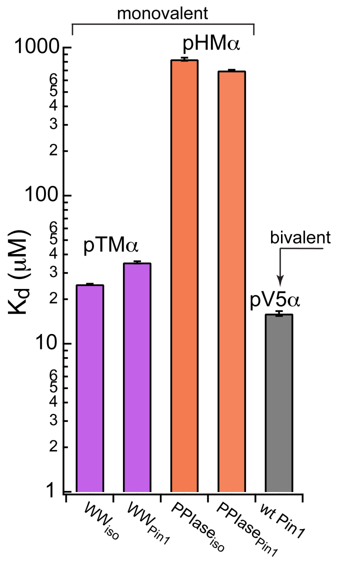

**Figure S11. Thermodynamic benefits of bivalent of Pin1-C-term PKCα interactions.** $K_d$ values for the monovalent interactions of the hydrophobic and turn motifs with isolated Pin1 domains and full-length Pin1 are contrasted with the $K_d$ value for the bivalent Pin1-pV5α interactions. ~3-fold enhancement for the pTM binding to the WW domain and ~60-fold enhancement of the pHM binding to the PPIase domain are attributed to bivalency. The $K_d$ values used for this plot were obtained in the NMR-detected binding experiments #3, 4, 6, 7, and 14 (**Table S2**).

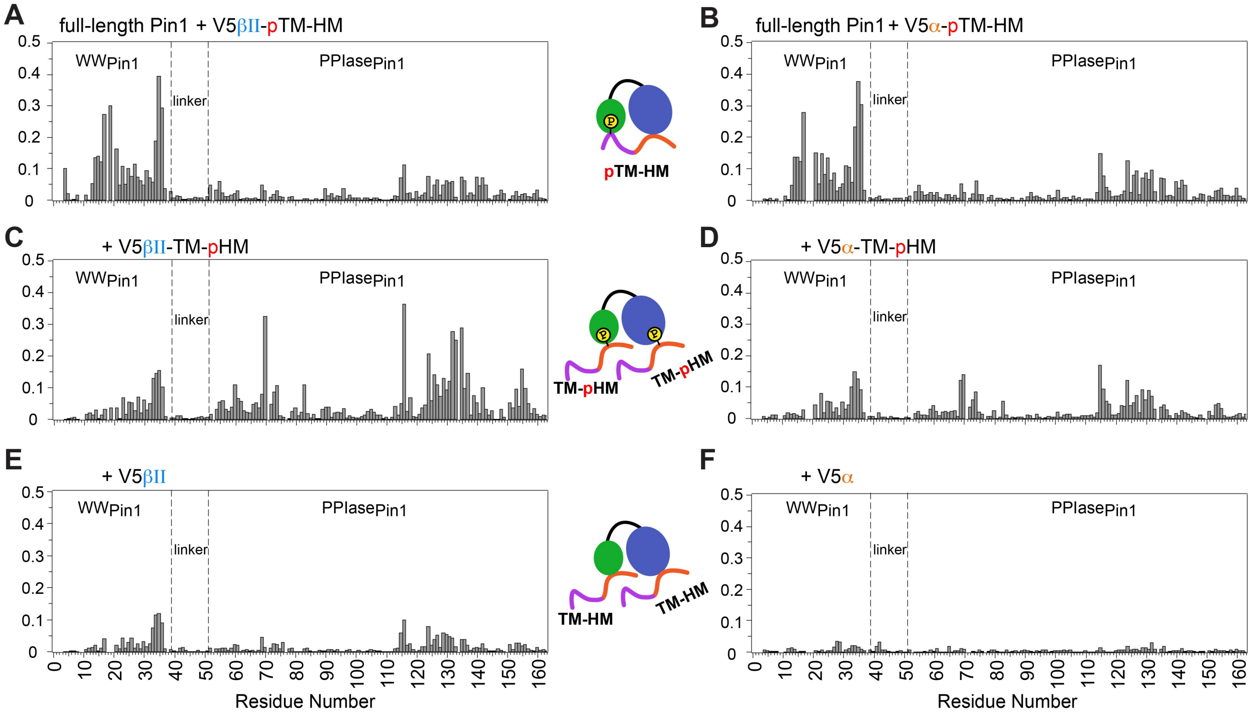

**Figure S12. Phosphorylation state of PKCβII/α C-term regions dictate the Pin1 binding mode.** Comparison of the Pin1 CSP plots of Pin1 obtained at maximum concentrations of V5βII-pTM-HM (**A**), V5α-pTM-HM (**B**), V5βII -TM-pHM (**C**), V5α-TM-pHM (**D**), V5βII (**E**), and V5α (**F**). Pin1 binds to the PKC C-term in a bivalent mode only when TM is phosphorylated (**A-B**). When TM is unphosphorylated, the hydrophobic motif interacts with both WW and PPIase domains (**C-F**). The protein concentration was 100 μM. Other details are given in **Table S2**.

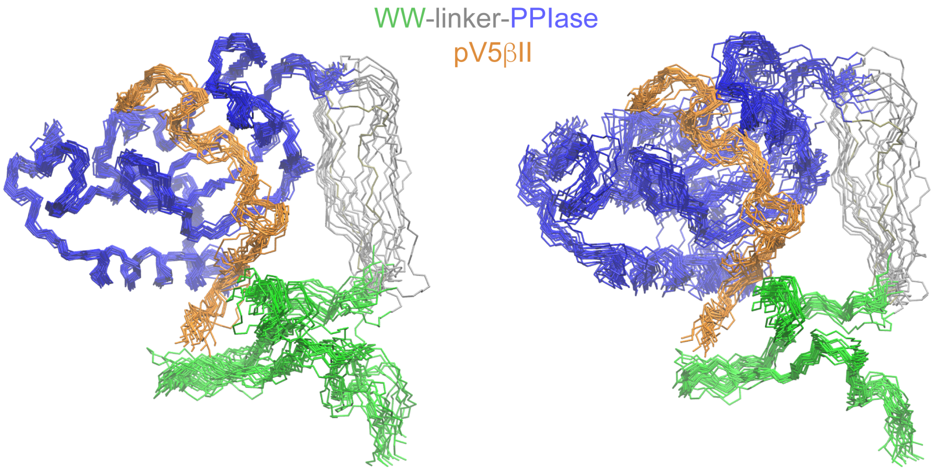

**Figure S13.** NMR ensemble of the Pin1::pV5βII complex (PDB ID 8SG2) reveals a novel Pin1 conformation. 20 lowest-energy structures of the Pin1::pV5βII ensemble superimposed using either the PPIase domain (left panel) or WW domain (right panel).

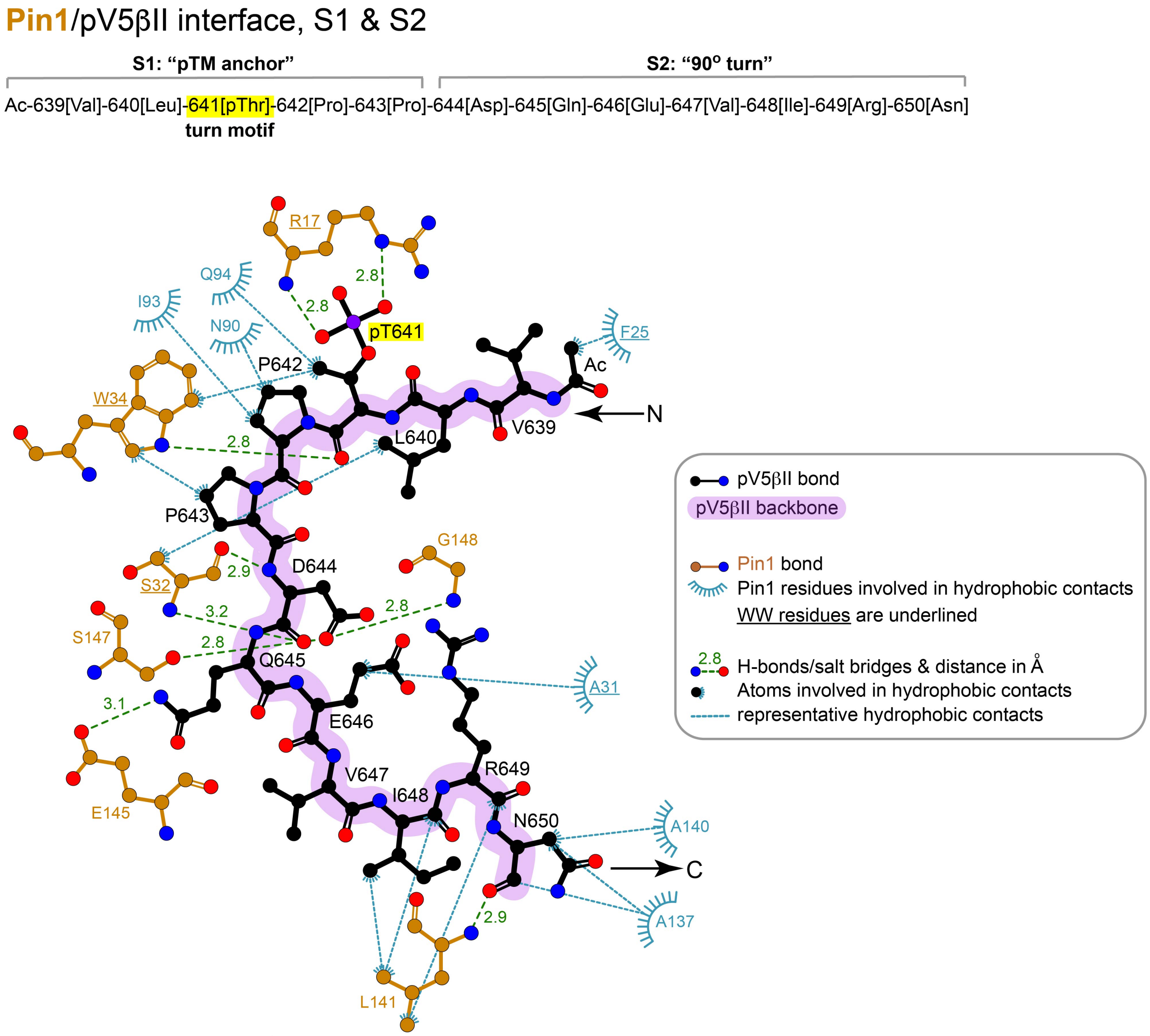

**Figure S14.** 2D LigPlot[+] diagram of representative Pin1 interactions with residues 639-650 ("pTM anchor" and "90° turn") of pV5βII. The lowest-energy structure of the Pin1::pV5βII complex was used to generate the diagram. The contact cutoff for hydrophobic contacts is 4.0 Å. The turn motif is highlighted in yellow.

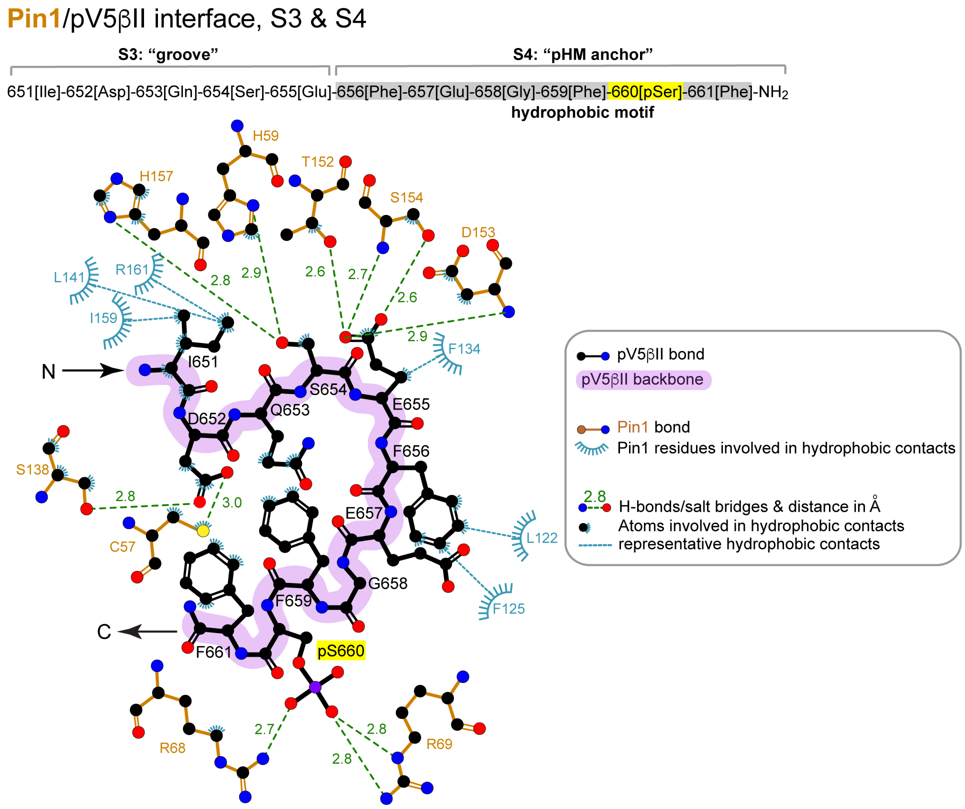

**Figure S15.** 2D LigPlot[+] diagram of representative Pin1 interactions with residues 651-661 (“groove” and “pHM anchor” segments) of pV5βII. The lowest-energy structure of the Pin1::pV5βII complex was used to generate the diagram. The contact cutoff for hydrophobic contacts is 4.0 Å. The hydrophobic motif is highlighted in gray.

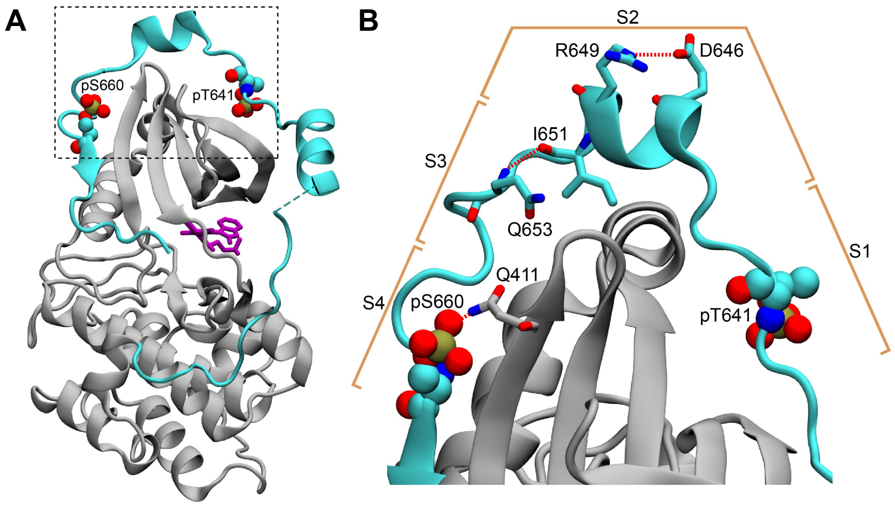

**Figure S16. The C-terminal tail in the structure of the PKCβII catalytic domain (PDB ID 2I0E). (A)** The C-terminal V5 domain (cyan) has elevated B-factors and peripherally interacts with the N-lobe of the catalytic domain (gray). **(B)** The intra-V5 R649-D646 salt bridge and the Q653(N-$H_N$)-(O=C)I651 H-bond that are also present in the Pin1-bound pV5βII are labeled. The S1-S4 segment notation that we used to analyze the Pin1::pV5βII complex is shown in the context of the catalytic domain structure.

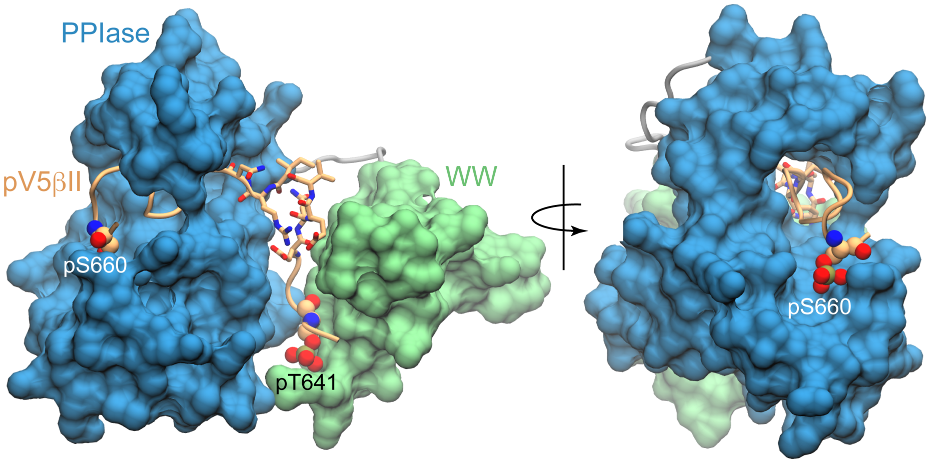

**Figure S17. The C-terminal part of pV5βII is threaded through the PPIase groove.** Space-filling representation showing the threading of pV5βII through the PPIase domain and its anchoring by the phosphate group of pS660. The "90° turn" segment is shown in licorice representation. The figures were prepared using the lowest-energy Pin1::pV5βII structure.

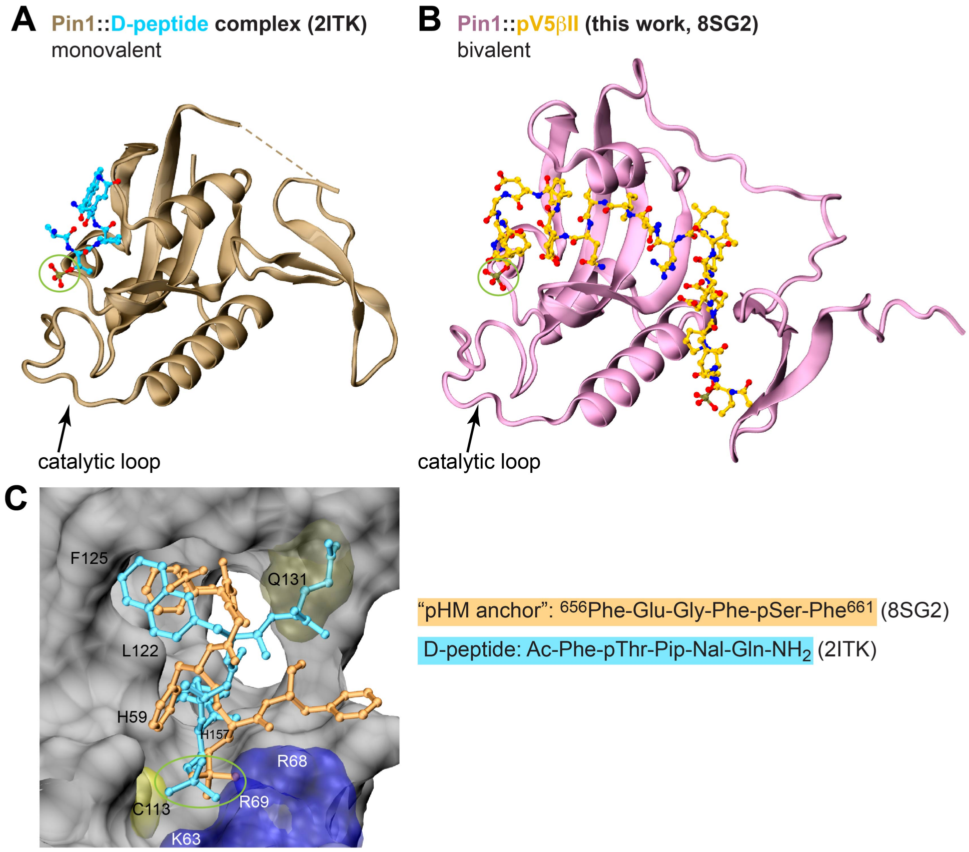

**Figure S18. Comparison of the binding poses between the D-peptide, a potent unnatural peptide inhibitor of Pin1, and the "pHM anchor" segment of pV5βII.** Crystal structure of the monovalent Pin1::D-peptide complex **(A)** and the lowest-energy NMR structure of the bivalent Pin1::pV5βII complex **(B)**. The phosphate group interacting with the catalytic loop is highlighted with a green circle. **(C)** The binding poses of the "pHM anchor" (dark yellow) and the D-peptide (cyan) in the catalytic site of Pin1. The Gln sidechain of the D-peptide occupies the space that is taken up by the Pin1 Gln131 sidechain in the Pin1::pV5βII complex.

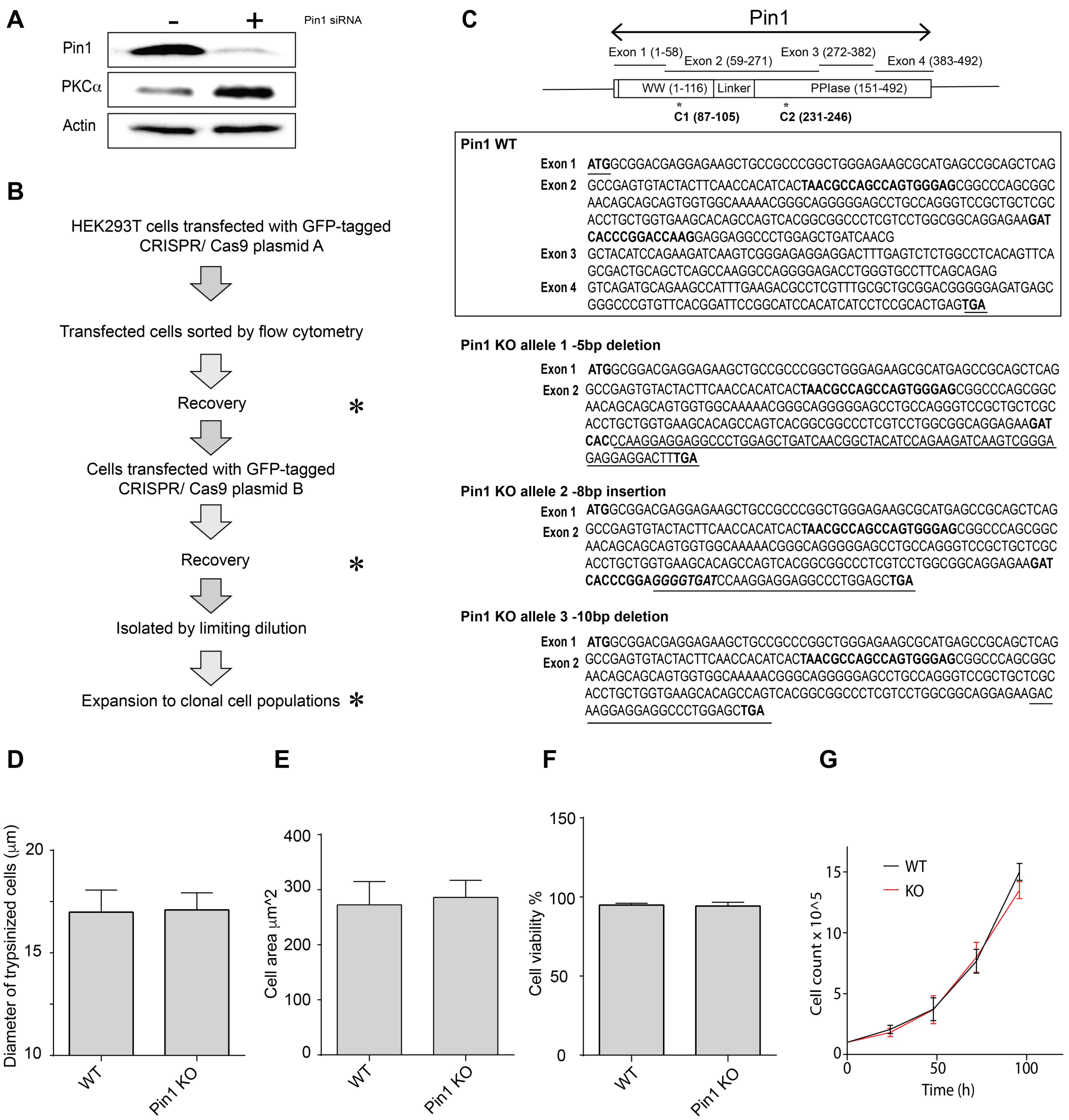

**Figure S19. PKCα levels in cells with reduced Pin1 function.** (**A**) HEK293T cells were transfected with mock or Pin1 siRNA as indicated at top, incubated for 72 hours in serum-replete medium, and cell lysates were prepared and analyzed by immunoblotting. Immunoblot profiles for Pin1, PKCα and actin are shown. Actin provides a normalizing signal. Pin1 knockdown (~90%) resulted in a significant elevation of PKCα levels. (**B**) Diagram outlining the workflow for generating clonally-derived Pin1 null HEK293T cells using CRISPR/Cas9. (**C**) At top is illustrated the Pin1 gene organization with codons present in each exon shown in parentheses. Domain organization of the Pin1 protein is shown at bottom. Middle panel shows the exon coding sequences for Pin1. The start and stop codons are highlighted (underlined, bold) as are the CRISPR/Cas9 targeting sequences in exon 2 (bold). Bottom panel shows the open reading frames of each of the three *Pin1* null alleles identified in clone C3. The natural start codon and the CRISPR/Cas9 targeting sequences in exon 2 are highlighted in bold. The short open reading frame extensions that lie downstream of each frameshift allele are underlined and the nonsense codons that terminate translation are indicated in bold. (**D**) The diameters (μm) of trypsinized wild-type or Pin1 KO HEK293T cells were determined using a Countess™ automated cell counter. (**E**) Cell areas ($\mu m^2$) of trypsinized wild-type or Pin1 KO HEK293T cells were determined by flow cytometry. (**F**) Viabilities of trypsinized wild-type or Pin1 KO HEK293T cells were assessed by trypan blue staining coupled to imaging with a Countess™ automated cell counter. (**G**) Growth rates of wild-type and Pin1 KO HEK293T cells were followed in a 96h window with an initial count of $1 \times 10^5$ cells (n = 3). Similar data showing no difference between wild-type and KO cell growth rates were also observed when the initial count was reduced to $5 \times 10^4$ cells (data not shown).

### TABLES

**Table S1.** Properties of the 18 PKC C-terminal-derived peptides used in this study. All peptides have acetylated N-termini and amidated C-termini. TM and HM stand for the turn and hydrophobic motifs, respectively. The phosphorylated Thr of the TM and phosphorylated Ser of the HM are shown in red. Peptides having “V5” in their name contain both TM and HM. Peptides starting with “p” indicate that the peptide is phosphorylated at either one or both motifs, the latter only for the “V5” peptides.

| Source | Peptide name | Sequence | pI | Solu-bility (water) | Vendor name |
|---|---|---|---|---|---|
| PKCα | 1. V5α | $^{632}$RGQPVLT$^{638}$PPDQLVIANIDQSDFEGFS$^{657}$YVN$^{660}$ | 3.3 | Poor | Eton-Bioscience |
| | 2. V5α-pTM-HM | $^{632}$RGQPVLpT$^{638}$PPDQLVIANIDQSDFEGFS$^{657}$YVN$^{660}$ | 3.3 | Poor | Eton-Bioscience |
| | 3. V5α-TM-pHM | $^{632}$RGQPVLT$^{638}$PPDQLVIANIDQSDFEGFpS$^{657}$YVN$^{660}$ | 3.3 | Poor | Eton-Bioscience |
| | 4. pV5α | $^{632}$RGQPVLpT$^{638}$PPDQLVIANIDQSDFEGFpS$^{657}$YVN$^{660}$ | 3.3 | Poor | Eton-Bioscience |
| | 5. pTMα | $^{632}$RGQPVLpT$^{638}$PPDQ$^{642}$ | 7.9 | Good | Eton-Bioscience |
| | 6. pHMα | $^{646}$ANIDQSDFEGFpS$^{657}$YVN$^{660}$ | 0 | Poor | Eton-Bioscience |
| PKCβII | 7. V5βII | $^{640}$LT$^{641}$PPDQEVIRNIDQSEFEGFS$^{660}$F$^{661}$ | 3.3 | Good | Eton-Bioscience |
| | 8. pV5βII | $^{640}$LpT$^{641}$PPDQEVIRNIDQSEFEGFpS$^{660}$F$^{661}$ | 3.3 | Good | Eton-Bioscience Thermo-Fisher Scientific |
| | 9. pTMβII | $^{640}$LpT$^{641}$PPDQEVIR$^{649}$ | 3.9 | Good | Eton-Bioscience |
| | 10. pHMβII | $^{650}$NIDQSEFEGFpS$^{660}$F$^{661}$ | 0 | Good | Eton-Bioscience |
| | 11. V5βII-TM-pHM | $^{640}$LT$^{641}$PPDQEVIRNIDQSEFEGFpS$^{660}$F$^{661}$ | 3.3 | Good | Eton-Bioscience |
| | 12. V5βII-pTM-HM | $^{640}$LpT$^{641}$PPDQEVIRNIDQSEFEGFS$^{660}$F$^{661}$ | 3.3 | Good | Thermo-Fisher Scientific |
| | 13. Ext-pV5bII | $^{639}$VLpT$^{641}$PPDQEVIRNIDQSEFEGFpS$^{660}$FVN$^{663}$ | 3.3 | Good | Thermo-Fisher Scientific |
| | 14. Labeled-pTMβII | $^{640}$LpT$^{641}$PPDQEVIR$^{649}$ ($^{13}$C,$^{15}$N-labeled) | 3.9 | Good | Sigma-Aldrich |
| | 15. Labeled-pHMβII | $^{650}$NIDQSEFEGFpS$^{660}$F$^{661}$ ($^{13}$C,$^{15}$N-labeled) | 0 | Good | Sigma-Aldrich |
| α, βII | 16. SP-1 | LpTPPD | 0 | Good | Eton-Bioscience |
| βI | 17. SP-2 | LpTPTD | 0 | Good | Eton-Bioscience |
| α, variant | 18. pTMα P640A | $^{632}$RGQPVLpT$^{638}$PADQ$^{642}$ | 7.9 | Good | Eton-Bioscience |

**Table S2.** List of binding experiments carried out in this study, with the corresponding values of the dissociation constants $K_d$ obtained from the chemical-shift binding curves and/or lineshape analysis.

| Binding Experiment | Protein | V5 region, Max conc. (μM) | Residues selected for analysis | $K_d$ (μM) Chem. shift | $K_d$ (μM) Lineshape analysis |
|---|---|---|---|---|---|
| 1 | Pin1 | pTMβII, 1000 | K13, R14, M15, S16, G20, R21, V22, Y23, Y24, F25, N26, I28, N30, A31, S32, Q33, W34, E35, R36 | 9.1 ± 0.4 | 9.6 ± 0.1 |
| 2 | WW | pTMβII, 1000 | W34, S16, E35, Q33, G20, K13, H27, M15, N26, R14, S32, Y24, V22, A31, F25, Y23, T29, W34ε, R21, N30, E12, I28 | 14.3 ± 0.2 | 16.0 ± 0.1 |
| 3 | Pin1 | pTMα, 1800 | R36, Q33, M15, K13, R21, N26, I28, F25, E35, G20, Y24, V22, E12, R14 | 35.4 ± 0.6 | 32.7 ± 0.1 |
| 4 | WW | pTMα, 1800 | S32, S16, A31, E35, N26, N30, Q33, M15, G20, R14, K13, Y24, Y23, W34, V22 | 25.2 ± 0.2 | 24.0 ± 0.1 |
| 5 | WW | pHMα, 1800 | Q33, S32, W34ε, W34, E35, F25, V22, S16, G20, Y24, A31, N30, Y23, T29 | 1006 ± 9 | |
| 6 | PPIase | pHMα, 1878 | A116, A124, D136, D153, E135, F125, F134, G123, H59, K117, K132, M130, Q129, Q131, S114, S115, S126, W73, R68, R69 | 835 ± 20 | |
| 7** | Pin1 | pHMα, 1800 | δH and δN: S18, G20, V22, Y23, Y24, F25, T29, N30, S32, Q33, W34, W34 ε, E35, S58, S72, S114, S115, A116, K117, G128, Q129, Q131, K132, D136, T152, D153;<br>δH only: Q66, R68, R69, G120, F125, M130, S154;<br>δN only: H59, Q66, R68, R69, W73, W73ε, K82, D121, A124, F125, S126, F134, E135, A31 | **WW:** 725 ± 9<br>**PPIase:** 701 ± 7 | |
| 8 | WW | pHMβII, 2550 | W34, S16, E35, Q33, G20, S32, Y24, V22, A31, F25, Y23, T29, N30, W34 ε | 782 ± 9 | 809 ± 4 |
| 9 | PPIase | pHMβII, 1125 | S65, I156, L61, V55, A124, K117, V150, L141, L60, R127, S138, T152, S72, E135, G155, Q129, H59, S126, G128, F125, D136, S154, K132, F134, R69, S115 | 133 ± 1 | 127 ± 1 |
| 10** | Pin1 | pHMβII, 1350 | δH and δN: G20, V22, Y24, F25, A31, S32, Q33, W34, E35, R54, V55, H59, Q66, R68, R69, S72, K82, S114, S115, K117, A124, F125, S126, R127, Q129, K132, F134, E135, D136, S138, F139, L141, V150, T152, I156;<br>δH only: T29, R36, L61, S65, W73, L122, G128;<br>δN only: N30, G123, G155 | **WW:** 257 ± 9<br>**PPIase:** 75.0 ± 1.1 | |
| 11 | Pin1 | V5βII-pTM-HM, 460 | W34, S18, E35, S16, Q33, G20, R14, K13, M15, N26, V22, Y24, N30, F25, A31, I28, S32, E12, Y23, T29, R21, R36, S115, F134, F139, L141, E51, R54, R142, A140, A53, V55, I158, S114, S126, Q129, E135, Q131, M130, F125, G128, R69, T152, K117 | 21.0 ± 0.5 | 20.1 ± 0.1 |
| 12** | Pin1 | V5α-TM-pHM, 1320 | δH and δN: V22, Y24, A31, S32, W34, E35, R69, S72, S114, S115, A116, K117, F125, G128, M130, Q131, L141, T152, D153;<br>δH only: G20, F25, T29, H59, W73, K82, S126, F134, E135; | **WW:** 688 ± 21<br>**PPIase:** 757 ± 16 | |

| Binding Experiment | Protein | V5 region, Max conc. (μM) | Residues selected for analysis | $K_d$ (μM) Chem. shift | $K_d$ (μM) Lineshape analysis |
|---|---|---|---|---|---|
| | | | δN only: Q33, S55, R68, G123, Q129, S154 | | |
| 13** | Pin1 | V5βII-TM-pHM, 1640 | δH and δN: G20, V22, Y24, F25, A31, S32, Q33, W34, V55, H59, Q66, S72, K82, S114, S115, K117, A124, F125, S126, R127, Q129, K132, F134, E135, D136, S138, F139, L141, V150, T152, I156; δH only: T29, E35, R36, R54, L61, S65, R68, R69, W73, L122, G128; δN only: N30, R54, R68, R69, L122, G123, G155 | **WW:** 213 ± 7 **PPIase:** 93.6 ± 1.5 | |
| 14 | Pin1 | pV5α, 1530 | A31, F25, G20, K13, M15, N26, N30, Q33, R14, R21, S32, T29, V22, Y23, Y24, A116, E135, E51, F125, F134, F139, G123, G128, K117, K82, L141, M130, Q129, Q131, R68, R69, S114, S115, S126, S71, S72, T152, W73ε, W73 | 16.0 ± 0.6 | 12.0 ± 0.2 |
| 15 | Pin1 | pV5βII, 400 | M15, V22, F25, N26, A31, S32, R54, V55, S72, S115, K117, G128, Q129, D136, F139, L141, S147, V150, T152, S154, G155, R69, S115, K132, F125, S126 | N/D | 1.5 ± 0.1 |
| 16 | Pin1 | V5βII, 1650 | K13, S16, G20, V22, T29, A31, S32, Q33, W34, E35, R68, W73, K112, S114, S115, A116, D121, G123, A124, F125, S126, G128, Q129, M130, Q131, E135, D136, L141, T152, G155 | 964 ± 34 | |
| 17 | Pin1 | V5α, 500 | | N/A | |
| 18 | Pin1, pTMβII-complexed (98% saturated) | pHMβII, 500 | S65, V55, A124, K117, V150, L141, L60, S138, T152, S72, E135, G155, Q129, H59, S126, G128, F125, D136, S154, K132, F134, R69, S115, G123, A107, Q131, S114 | 117 ± 2 | |
| 19 | Pin1, C113S variant | pV5βII, 350 | E12, R14, M15, F25, N26, T29, A31, R54, H64, S72, D112, A116, F125, S126, G128, Q129, M130, F134, A137, F139, L141, S147 | N/D | 3.4 ± 0.1 |

**2-site binding model
N/D: Not determined; affinity is too high for the chemical-shift based analysis.
N/A: Impossible to determine the $K_d$ due to very low affinity.

**Table S3.** List of the NMR samples and experiments for the structure determination of the complex. Sample 2* was prepared in the buffer containing 100% $D_2O$.

| ID | Complex | pV5βII conc. (mM) | Pin1 conc. (mM) | Bound pV5βII, % | Bound Pin1, % | Experiments |
|---|---|---|---|---|---|---|
| 1 | [U-$^{13}$C, $^{15}$N] Pin1 + pV5βII | 1.05 | 0.9 | 98 | 84 | --2D experiments: [$^{15}$N, $^{1}$H] HSQC, ct [$^{13}$C, $^{1}$H] HSQC, (HB)CB(CGCDCE)HE, (HB)CB(CGCD)HD<br><br>--3D experiments for backbone assignment: HNCACB, CBCA(CO)NH, HNCO, HN(CA)CO<br><br>--3D experiments for side-chain assignments: C(CO)NH, H(CCO)NH, HNHA, HNHB<br><br>--NOESY experiments: 3D $^{15}$N-edited NOESY-HSQC, 3D [F1] $^{13}$C,$^{15}$N-filtered NOESY-$^{15}$N-HSQC |
| 2* | [U-$^{13}$C, $^{15}$N] Pin1 + pV5βII | 1.05 | 0.9 | 98 | 84 | --$^{1}$H/$^{2}$D exchange<br><br>--2D ct [$^{13}$C, $^{1}$H] HSQC, 2D ct $^{1}$H-$^{13}$C aromatic HSQC<br><br>--3D experiments for sidechain assignments: HCCH-COSY, HCCH-TOCSY<br><br>--NOESY experiments: 3D $^{13}$C-edited NOESY-HSQC, 3D $C^{aro}$-edited NOESY-HSQC, 3D [F1] $^{13}$C,$^{15}$N-filtered NOESY-$^{13}$C-HSQC |
| 3 | [U-$^{13}$C, $^{15}$N] Pin1 + pV5βII | 1.0 | 1.3 | 76 | 99 | --2D [F1] $^{13}$C,$^{15}$N-filtered NOESY<br>--2D [F2] $^{13}$C,$^{15}$N-filtered NOESY<br>--2D [F1, F2] $^{13}$C,$^{15}$N-filtered NOESY<br>--2D [F1, F2] $^{13}$C,$^{15}$N-filtered TOCSY |
| 4 | [U-$^{13}$C, $^{15}$N] PPIase + pHMβII | 0.8 | 5.5 | 97 | 14 | --3D [F1] $^{13}$C, $^{15}$N-filtered NOESY-$^{15}$N-HSQC<br>--3D [F1] $^{13}$C, $^{15}$N-filtered NOESY-$^{13}$C-HSQC |
| 5 | [U-$^{13}$C, $^{15}$N] PPIase + pHMβII | 0.5 | 1.8 | 25 | 91 | --2D [F1] $^{13}$C,$^{15}$N-filtered NOESY<br>--2D [F2] $^{13}$C,$^{15}$N-filtered NOESY<br>--2D [F1, F2] $^{13}$C,$^{15}$N-filtered NOESY<br>--2D [F1, F2] $^{13}$C,$^{15}$N-filtered TOCSY |
| 6 | PPIase + [U-$^{13}$C,$^{15}$N Phe] pHMβII *peptide ID #15 (Table S1)* | 0.5 | 2.5 | 19 | 93 | --3D [F1] $^{13}$C,$^{15}$N-filtered NOESY-$^{15}$N-HSQC<br>--3D [F1] $^{13}$C,$^{15}$N-filtered NOESY-$^{13}C^{aro}$-HSQC<br>--3D [F1] $^{13}$C,$^{15}$N-filtered NOESY-$^{13}C^{ali}$-HSQC |

| ID | Complex | pV5βII conc. (mM) | Pin1 conc. (mM) | Bound pV5βII, % | Bound Pin1, % | Experiments |
|---|---|---|---|---|---|---|
| 7 | PPIase + [U-$^{13}C$,$^{15}N$ Pro, Ile, Val] pTMβII *peptide ID #14 (Table S1)* | 1.0 | 1.5 | 66 | 98 | --3D [F1] $^{13}C$,$^{15}N$-filtered NOESY-$^{15}N$-HSQC<br>--3D [F1] $^{13}C$,$^{15}N$-filtered NOESY-$^{13}C$-HSQC |
| 8 | [U-$^{13}C$, $^{15}N$] Pin1 + Ext-pV5βII *peptide ID #13 (Table S1)* | 1.3 | 1.0 | >95% | | --3D [F1] $^{13}C$,$^{15}N$-filtered NOESY-$^{15}N$-HSQC<br>--3D [F1] $^{13}C$,$^{15}N$-filtered NOESY-$^{13}C$-HSQC |
| 9 | [U-$^{13}C$, $^{15}N$] Pin1 + Ext-pV5βII *peptide ID #13 (Table S1)* | 0.8 | 1.3 | | >95% | --2D [F1] $^{13}C$,$^{15}N$-filtered NOESY<br>--2D [F2] $^{13}C$,$^{15}N$-filtered NOESY<br>--2D [F1, F2] $^{13}C$,$^{15}N$-filtered NOESY<br>--2D [F1, F2] $^{13}C$,$^{15}N$-filtered TOCSY |

**Table S4.** NMR restraints statistics for the CYANA structure calculation.

| | |
|---|---|
| **No. of NOE restraints for Pin1** | |
| No. of NOE-based distance restraints for Pin1 | |
| Total | 2780 |
| Intra-residue (\|i – j\| = 0) | 1071 |
| Sequential (\|i – j\| = 1) | 624 |
| Medium-range (1< \|i – j\| ≤ 4) | 339 |
| Long-range (\|i – j\| ≥ 5) | 662 |
| NOE restraints per residue | ~17 |
| No. of hydrogen bond restraints | 84 |
| No. of dihedral angle restraints | 244 |
| Total no. of restraints | 3024 |
| Total no. of restraints per residue | ~18.6 |
| **No. of NOE restraints for pV5βII** | |
| No. of NOE-based distance restraints for Pin1 | |
| Total | 241 |
| Intra-residue (\|i – j\| = 0) | 147 |
| Sequential (\|i – j\| = 1) | 67 |
| Medium-range (1< \|i – j\| ≤ 4) | 24 |
| Long-range (\|i – j\| ≥ 5) | 3 |
| NOE restraints per residue | ~11.0 |
| No. of dihedral angle restraints | 0 |
| Total no. of restraints | 241 |
| Total no. of restraints per residue | 11 |
| Total No. of intermolecular NOEs = 75 | |
| Total no. of structures calculated | 500 |
| Total no. of structures refined | 50 |
| No. of structures used | 20 |
| No. of restraint violations | |
| Dihedral angle of > 5° | |
| Distance of > 0.2 Å | |
| Van der Waals | 149 |
| Ramachandaran plot (Procheck) | |
| Most favoured regions (%) | 89.3 |
| Additionally allowed regions (%) | 10.7 |
| Generously allowed regions (%) | 0.0 |
| Total allowed regions (%) | 100.0 |
| Disallowed regions (%) | 0.0 |
| Ramachandaran plot (Molprobity) | |
| Most favoured regions (%) | 96.2 |
| Allowed regions (%) | 3.8 |
| Total allowed regions (%) | 100.0 |
| Disallowed regions (%) | 0.0 |
| RMSD from mean structure coordinates (Å) | |
| Backbone | 0.9 |
| Average heavy atom | 1.2 |
